# Automated Monitoring of Insects system: Building a non-lethal and scalable solution for monitoring of nocturnal insects

**DOI:** 10.64898/2026.02.06.704388

**Authors:** William Lord, Simon Teagle, Joshua Alton, Gabriel Bannister, Jonas Beuchert, Kim Bjerge, Dylan Carbone, Alba Gomez Segura, Tim Howson, Toke Thomas Høye, Jenna Lawson, Abhi Ravivarma, David B. Roy, Dan Rylett, Grace Skinner, Alan Warwick, Tom August

## Abstract

There is growing evidence that human-induced climate change and habitat loss are having negative impacts on insect populations. New technologies have a vital role in improving and expanding global biodiversity monitoring capacity to understand where change is happening and to support restoration.

Monitoring of insects traditionally needs entomologists in the field, but insect camera traps powered by AI are emerging as a scalable approach to monitoring semi-autonomously. These systems attract, detect, and identify insects using AI algorithms and are being developed by a network of researchers across the world, notably in Europe and North America.

The first version of a system for monitoring nocturnal insects was developed by Bjerge et al. 2021. An Automated Light Trap to Monitor Moths (Lepidoptera) Using Computer Vision-Based Tracking and Deep Learning. Here we describe the second generation of the system as an open-source solution. This paper aims to enable anyone to build their own system, and to iterate and improve the design for their needs. This system captures images at set intervals or based on motion detection to monitor insects that are attracted to lights at night. The UKCEH Automated Monitoring of Insects System (UKCEH AMI-system) is an insect camera trap designed using a single board computer, USB camera and attractant lights as the primary components along with peripheral accessories to make an autonomous system capable of long-term deployment in the field.

**Graphical Abstract:** 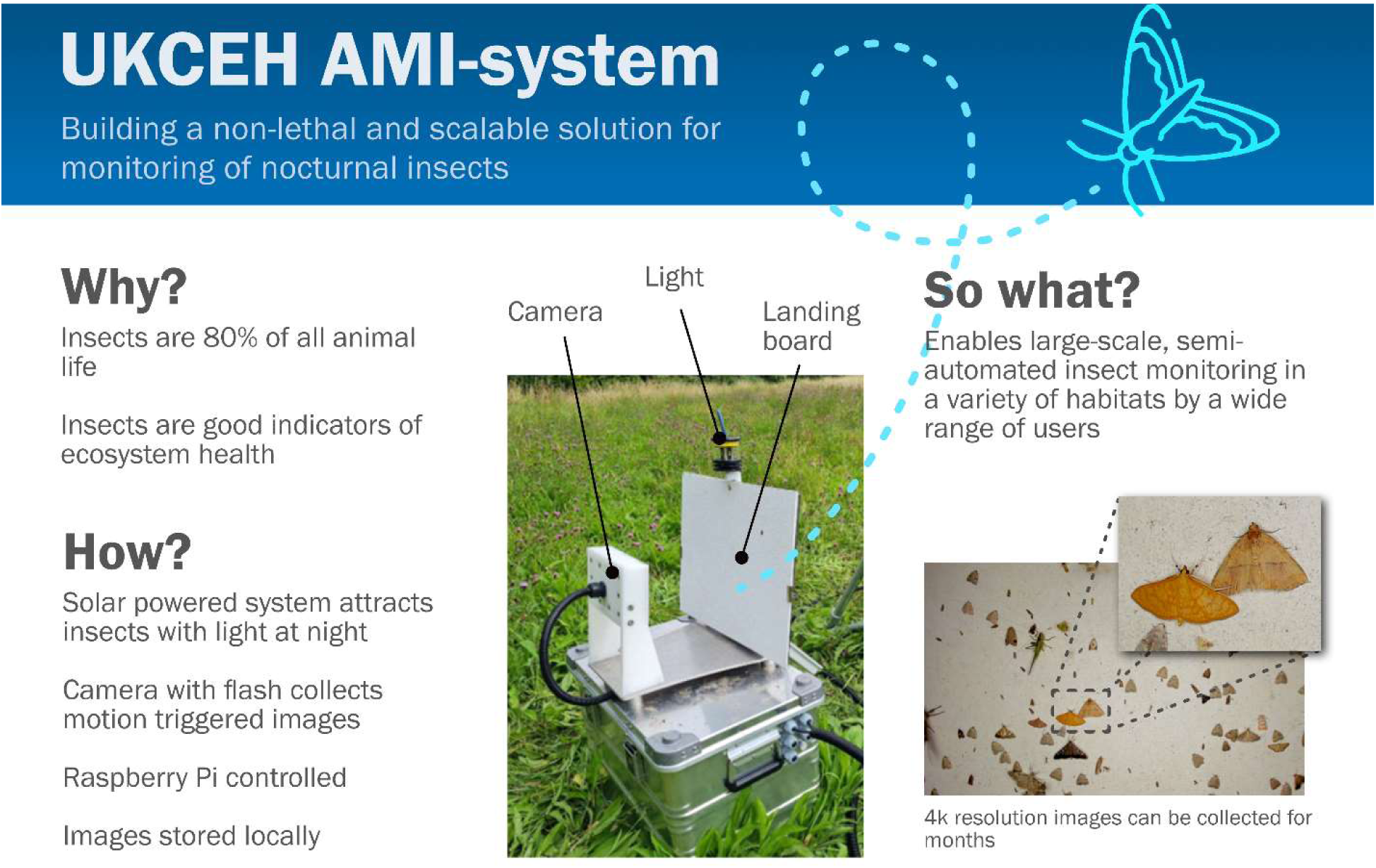

## 1. Hardware in context

There is growing evidence of widespread declines in insect diversity and abundance [1], caused by a range of environmental pressures such as habitat loss and climate change [2]. Robust monitoring data for insects is a priority for action [3], and necessary to identify solutions that lead to nature conservation, alongside economic prosperity. Insects are a critical component of biodiversity, playing a vital role in all terrestrial ecosystems, comprising approximately 80% of animal life [4] and providing critical ecosystem services such as pollination, pest control, nutrient cycling, and culture services. The huge diversity of insects and their rapid response to environmental conditions also makes them highly effective as indicators of wider biodiversity condition.

The solution we have developed, and field tested, is the UKCEH Automated Monitoring of Insects System (UKCEH AMI-system). This builds on a system for monitoring nocturnal insects first published by Bjerge et al. [5]. We focus on a range of insects attracted to light (i.e. moths, and a range of other species groups), but note that spiders, molluscs and other invertebrates have been recorded by the traps as well. The system uses a light and camera to capture images. These images can then be classified using Artificial Intelligence (AI) algorithms and manually verified with entomologists using platforms such as ANTENNA (antenna.insectai.org). The system is deployed with either power by mains/grid electricity using a 12 V DC adapter or through a solar powered charging setup of 12 V batteries.

The field of automated monitoring of insects with camera systems is relatively new, however, a number of systems, open and closed design, are already available. Notably the commercially available Diopsis system is capable of monitoring nocturnal and day-flying insects [6]. Other systems include the Moth Scanner which uses digital SLR camera to capture images [7,8], the MothBox which offers an open design [9], AutoMoth which uses a smartphone and bespoke app for capturing images [10], and others. In contrast to these systems, the UKCEH AMI-system aims to maximise the duration of deployment and weatherproof/ruggedness of the system, albeit at increased costs.

UKCEH AMI-systems, totallng 194, are now in use in over 32 countries across Asia, the Americas, Africa and Europe, supporting a range of projects. UKCEH AMI-systems have been deployed in arctic, temperate, and tropical systems, in open grassland and closed forest, and have endured tropical rainstorms as well as extreme heat. Users of the UKCEH AMI-system are diverse, and their questions similarly varied. This includes policy makers looking for tools to monitor the impact of policy decisions, businesses interested in evidencing their biodiversity action (e.g. habitat creation), researchers monitoring invasive species, farmers investigating sustainable farming practices, and financial institutions looking for methods to measure biodiversity credits.

## 2. Hardware description

The UKCEH AMI-system described here is a tool for researchers to study nocturnal insects attracted to light. With the capability of programmable intervals for autonomous data capture, weatherproof design and large storage capacity, the system can be deployed for long durations through the field season.

Using experience from field deployments we enhanced the original design of an insect camera system [5], to simplify the design for manufacturing and assembly, make operation more intuitive for users, improve image quality, ensure robustness and weather proofing, and support autonomous field usage for long-term deployments (figure 1).

**Figure 1.**
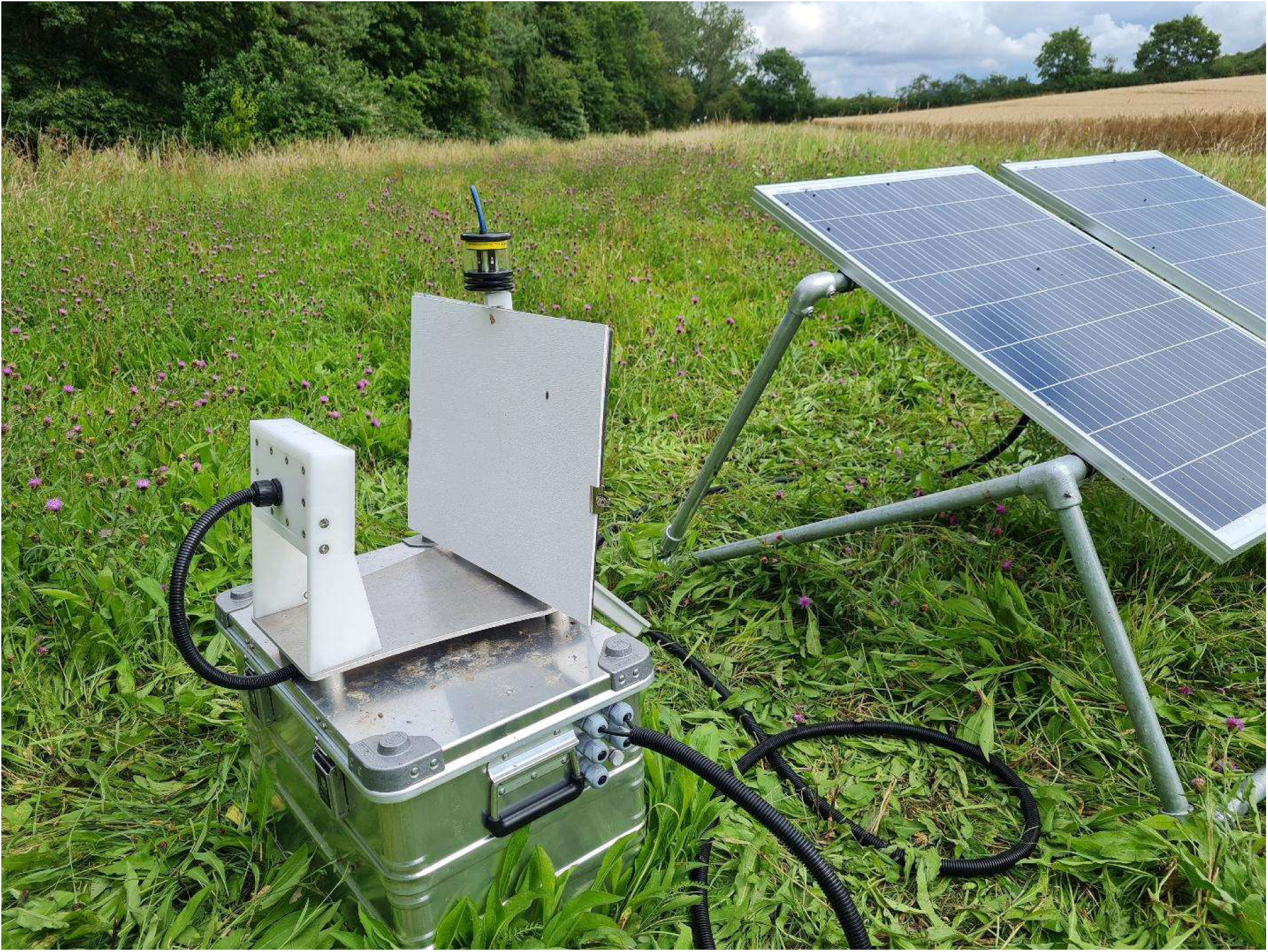
The UKCEH AMI-system consists of a solar panels (right), and a battery box (bottom), on top of which is a camera housing (left), a white landing board (right) - where insects land - and an attractant light (on top of the backboard). The electronics enclosure is mounted to the back of the backboard (not visible).

To achieve these improvements, we reduced the number of electronic components, ensured the system is protected against power interruption, can be turned off for periods of time to reduce power consumption, reinforced weatherproofing with an IP67 design, added solar charging capability for long term deployments in remote areas, and improved the LED light ring. Along with all these, we also paid attention to the environmental impact of material used in construction, the commercial availability of sub-components and use a modular design to facilitate field-based repairs and maintenance, and future technical developments.

The majority of the components that make up the UKCEH AMI-system are contained within the electronics enclosure. All the electronics components used for the operation of the system are described below and listed in the Bill of Materials.

We chose a Raspberry Pi model 4B for a single board computer to maintain and control the system operation. This model was chosen to allow for the ability to interface a high-resolution USB 3.0 video camera with high bandwidth data storage capability, along with sufficient processing power to manage the load during increased levels of activity (rapid image count and data storage). This choice also allows for future developments using different cameras and exploring the application of AI models on the device for processing images.

For the digital camera, we chose a Logitech BRIO 4K web camera with USB 3.0 to capture images of insects. The camera provides high resolution images (4K) and is compatible with the open source *motion* software [11] that enables easy camera configuration, automated operation using scripts, and motion detection.

We use a Super Watchdog Hat from Sequent Microsystems on the Raspberry Pi. Besides a watchdog timer, it provides an uninterrupted power supply (UPS) for the Raspberry Pi. In case of an unexpected power loss, it supplies power from an 18650 lithium-ion battery until pending tasks are completed and the system has safely shut down. This prevents corruption of data, i.e. the pictures being collected by the system.

To maintain the clock when the system is powered off we use the Adafruit RTC DS3231, a real-time clock, with a CR1220 coin cell battery. This maintains the clock settings, crucial metadata information embedded into the images captured for analysis.

There are two data storage devices in the system, a micro-SD card in the Raspberry Pi, and a solid-state drive (SSD) attached to the Raspberry Pi for storing pictures. A Raspberry Pi OS image is flashed onto the micro-SD card of 64 GB capacity, this contains all the software that is operating the UKCEH AMI-system. The SSD is a high capacity and fast read/write capable storage utility used to store all the pictures captured by the system. This is also a USB 3.0 device that can be swapped during data retrieval to speed up data collection.

The 12 V timer relay serves 2 functions, first to reduce power consumption and second to control the operating regime for the UKCEH AMI-system. By introducing the 12 V timer relay we are able to fully shutdown the system, meaning no power requirements during sleep intervals, with the timer relay much more energy efficient in these times. When scheduled to switch on, the relay boots up the system, this is useful for field deployments to operate the system autonomously during nocturnal periods. This also permits scheduling nights when the system does not run, limiting any negative impacts on local insect populations.

An LED light ring fitted around the camera housing supports illumination for the photography, these are a bespoke design made using 18 LED’s and 6 resistors mounted on to a PCB. The PCB construction is single sided FR4 (150 deg C) middle Tg, 1.2 mm thick with an immersion silver finish, size 170 mm x 90 mm. See Zenodo repository [12] for the bill of materials and Gerber files.

To stabilise the 12 V supply to the LED lighting (preventing inconsistencies in brightness), we use a 12 V DC-DC regulator, which prevents fluctuations in the voltage supplied to the LED light ring for photography hence maintains uniform lighting of pictures.

For the attractant light, used to draw the insects towards the camera, we chose the LepiLED [13] with a threaded bar, after iterations of designing our own UV light tube and evaluating other light options. LepiLED models are built using LEDs that emit UV, blue and green light in the electromagnetic spectrum (the three sensitivity peaks of most nocturnal insects) by eight Nichia Power LEDs contained in a weatherproof housing with an effective heat dissipation design. New models of attractant light are becoming available (e.g. EntoLED2 [14], entolight [15], MothBeam [16]) and the UKCEH AMI-system can work with any 12 V attractant light as long as there is sufficient battery capacity to operate it continuously. Light sources with a different voltage can be used in conjunction with a regulator.

The back board is manufactured from an aluminium composite panel. This allows the electronics enclosure and LepiLED to be securely mounted to the backboard as well as providing support to the landing board, which is the surface on which insects land to be photographed.

The landing board is a 5.5 mm thick 480 mm x 360 mm weather and boil proof (WBP) ply sheet which is painted with sandtex textured paint. The textured paint provides the insects with an easier surface to land on rather than the smooth surface of the backboard. In addition to this, the paint provides further protection from the weather and diffuses the light reflected off the surface, reducing glare. The landing board is secured to the back board using bulldog clips to prevent it from moving. Over a field season the landing board can become dirty, and while this does not significantly affect the performance of the device, we recommend cleaning or repainting boards seasonally.

The base plate joins the backboard, camera housing and battery box and is manufactured from 6082 aluminium. This material has excellent corrosion resistance as well as the strength required to maintain rigidity. We used a water jet cutter to make this part as it ensures high accuracy and is cost-effective, though other methods could be used. The high accuracy also ensures the camera housing and back board are in the same position each time the system is assembled, reducing the chance of images becoming out of focus.

The camera housing has the BRIO camera and LED light ring secured inside. The main body of the camera housing and back cover are machined from white acetal which has the required properties for withstanding a wide variety of environmental conditions. The front cover is manufactured from 4 mm polycarbonate. The polycarbonate ensures protection to the electronics inside and will not degrade with UV exposure, whilst letting all the light from the LED’s through. The front and back of the camera housing is sealed with a 1.5 mm silicone gasket. This gasket provides the waterproof seal for the housing.

The battery box is a metal Zarges box (mfr. part no. 40564). It has a series of glands installed into the box to allow for the solar panels and the UKCEH AMI-system to be connected. Inside the box there is space for three batteries (though we often use two); we use 120 Ah AGM lead acid deep cycle / marine batteries in parallel (12 V), but other similar batteries will also work. The maximum size per battery is 328 mm x 175 mm x 237 mm (length x width x height), to ensure they fit within the box. The box also contains a solar regulator (Morningstar Sunsaver) where the batteries, solar panels and UKCEH AMI-system are connected. The solar regulator controls the charging and load to ensure the batteries are not over charged or deeply discharged, which can damage them.

## 3. Design files summary

| **Design filename** | **File type** | **Open source license** | **Location of the file** |
| --- | --- | --- | --- |
| MothLEDBoard_v2.zip<br>LED ring light<br>PCB design files | CAD files | CERN Open Hardware Licence Version 2 - Strongly Reciprocal | https://doi.org/10.5281/zenodo.17580189 |
| AMI_hardware_drawings.zip<br>Mechanical construction design files | Figures / drawings | CERN Open Hardware Licence Version 2 - Strongly Reciprocal | https://doi.org/10.5281/zenodo.17580189 |
| AMI_solidworks_assembly_and_parts.zip<br>Solidworks files | Solidworks assembly and parts | CERN Open Hardware Licence Version 2 - Strongly Reciprocal | https://doi.org/10.5281/zenodo.17580189 |
| AMI_gasket_designs.zip<br>Design files for three gaskets used in the assemble of the camera enclosure | DXF files | CERN Open Hardware Licence Version 2 - Strongly Reciprocal | https://doi.org/10.5281/zenodo.17580189 |

MothLEDBoard_v2.zip – This file contains all of the CAD files needed for making the LED board that acts as the ring light around the camera. This illuminates the nocturnal insects to ensure good image quality and to some degree acts to attract insects to land on the illuminated board.

AMI_hardware_drawings.zip – This file contains PDF drawings of all of the components that need to be cut. This includes detailed information on materials, dimensions and the size and positioning of holes

AMI_solidworks_assembly_and_parts.zip – This file contains all the original SolidWorks part files and the assembly file. These files allow future users to more easily make edits to the designs we created.

AMI_gasket_designs.zip – This file contains three DXF files, one each for each of the gaskets that are used in the camera enclosure (figure 2)

**Figure 2.**
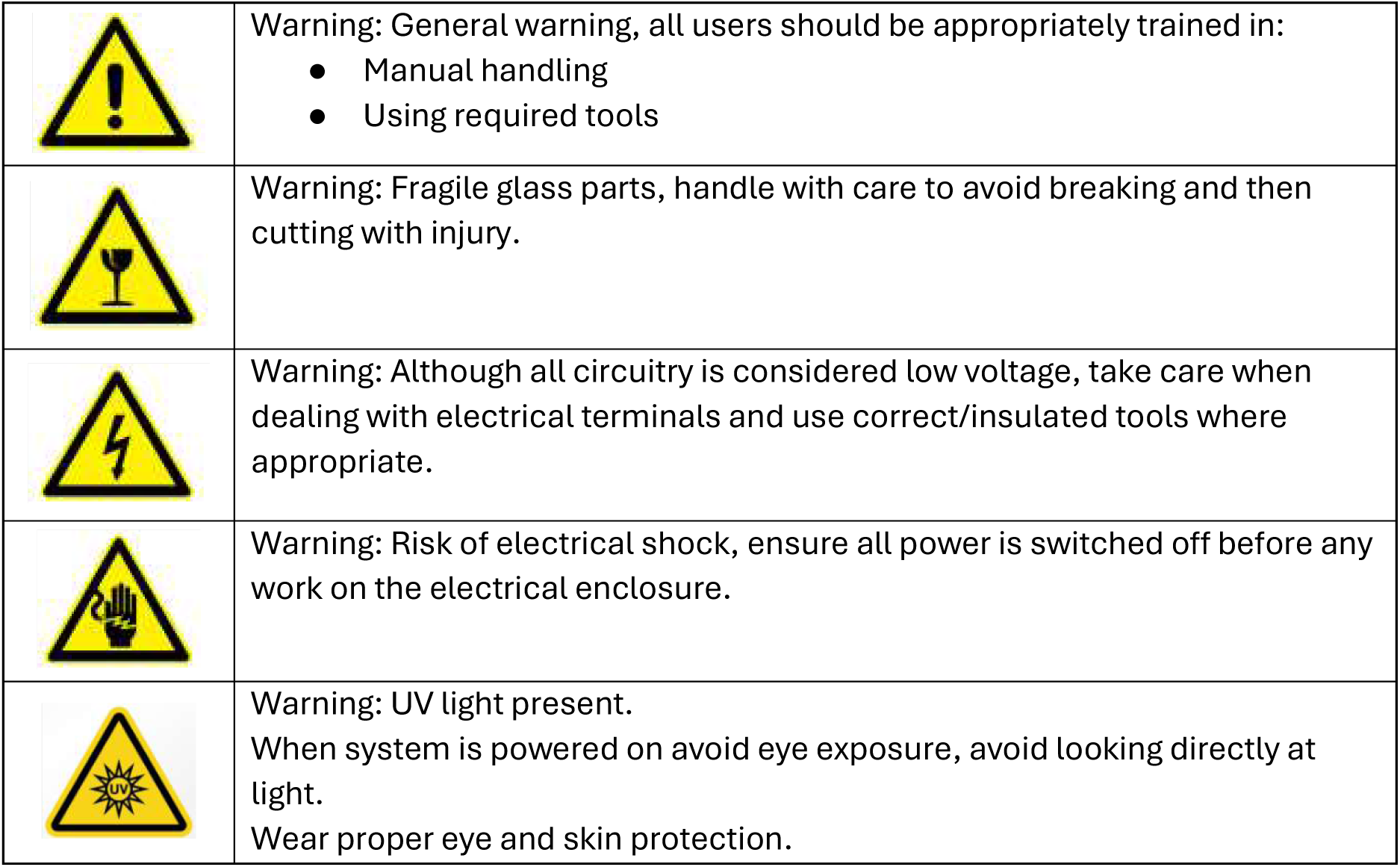
The above warnings should be displayed in materials disseminated for the construction of UKCEH AMI-systems, and displayed on the device once constructed, to ensure safe operation.

In addition to the design files listed above, the Zenodo repository also includes a detailed build manual, the bill of materials (BOM) and the image file for the Raspberry Pi [12].

### Bill of materials

The complete bill of materials is available with all other materials in our Zenodo repository for the UKCEH AMI-system [12]

## 5. Build instructions

The UKCEH AMI-system is broadly made up of five parts; the electronics enclosure, the battery box, the camera housing, the backboard, and the baseplate. The system has some hazardous components and so the warnings in figure 2 should be placed on the system, and in instructions provided with the system

### Assembling the camera enclosure

For more detailed step by step instructions, see the build manual the associated repository [12].

1. Mount the LED board (see files ‘MothLEDBoard_v2.zip’) into the front of the LED light and web camera housing using the 4 x M3 standoffs and 4 x m3 nylon screws (figure 3).
2. Outer and inner gaskets are cut and punched by hand from laser cut stencil using the AMOS outer and inner gasket dxf files.
3. Camera box is CNC milled.
4. Line the screw holes of the inner and outer gaskets with the holes on the housing and secure the LED cover using 28, M3 x 8 mm screws.
5. Turn the camera housing over and place the webcam into the slot and secure it with the 2 camera fixings.
6. Line the holes up in the back gasket with the holes in the web camera back and secure in place with the webcam back and 10 M5 x 12 mm screws.
7. Finally attach the left and right MK2 legs to the camera housing using 4, M6 x 20 mm screws.
8. Solder the 2-core cable to the positive and negative cable on the LED PCB board. Then pass the cable through the hole into the back of the camera housing. Next thread the two core cable and the USB cable from the camera through the cable gland and conduit.

**Figure 3.**
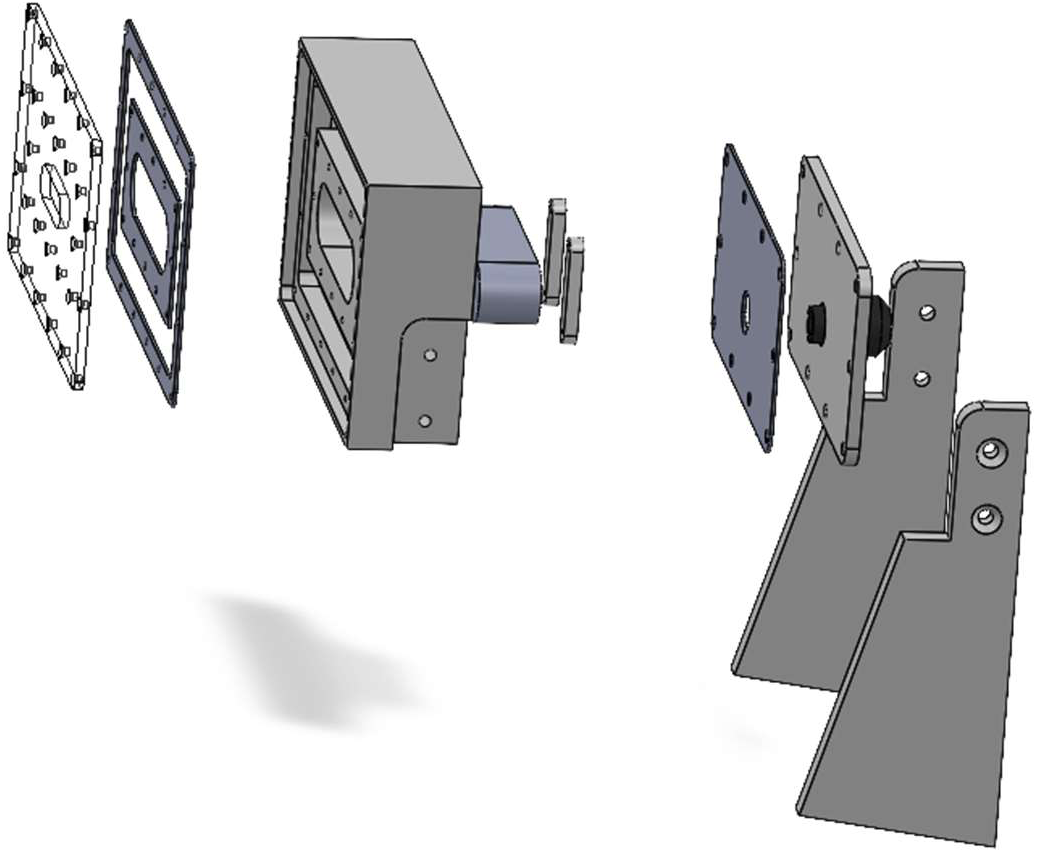
The assembly of the camera enclosure, including LED light ring, and the web camera.

### Assembling the battery box

The battery box is adapted from a Zarges box 40564 (410 x 600 x 400 mm).

1. Mark out the top of the battery box and drill the 5.3 mm diameter holes (figure 4). Use the battery box jig drawing for dimensions [12].
2. Mark out the hole position shown below and drill a 16 mm hole in the back left of the box to allow a power supply cable that will run from the battery box to the electronics enclosure (figure 5).
3. Install a solar regulator to the back of the battery box close to the top to allow for cables to connect.
4. Install fuses to the back right of the battery box, in line with the top of the box to allow space for cables to connect.
5. Wire the solar regulator to fuses following the manufacturers instructions
6. Pass the load wires from the fuses into conduit (300 mm long) securely attached to the hole made in step 2, ready to connect to the electronics enclosure.
7. For advice on batteries and solar panels see the section ‘Solar panel and battery recommendations’

**Figure 4:**
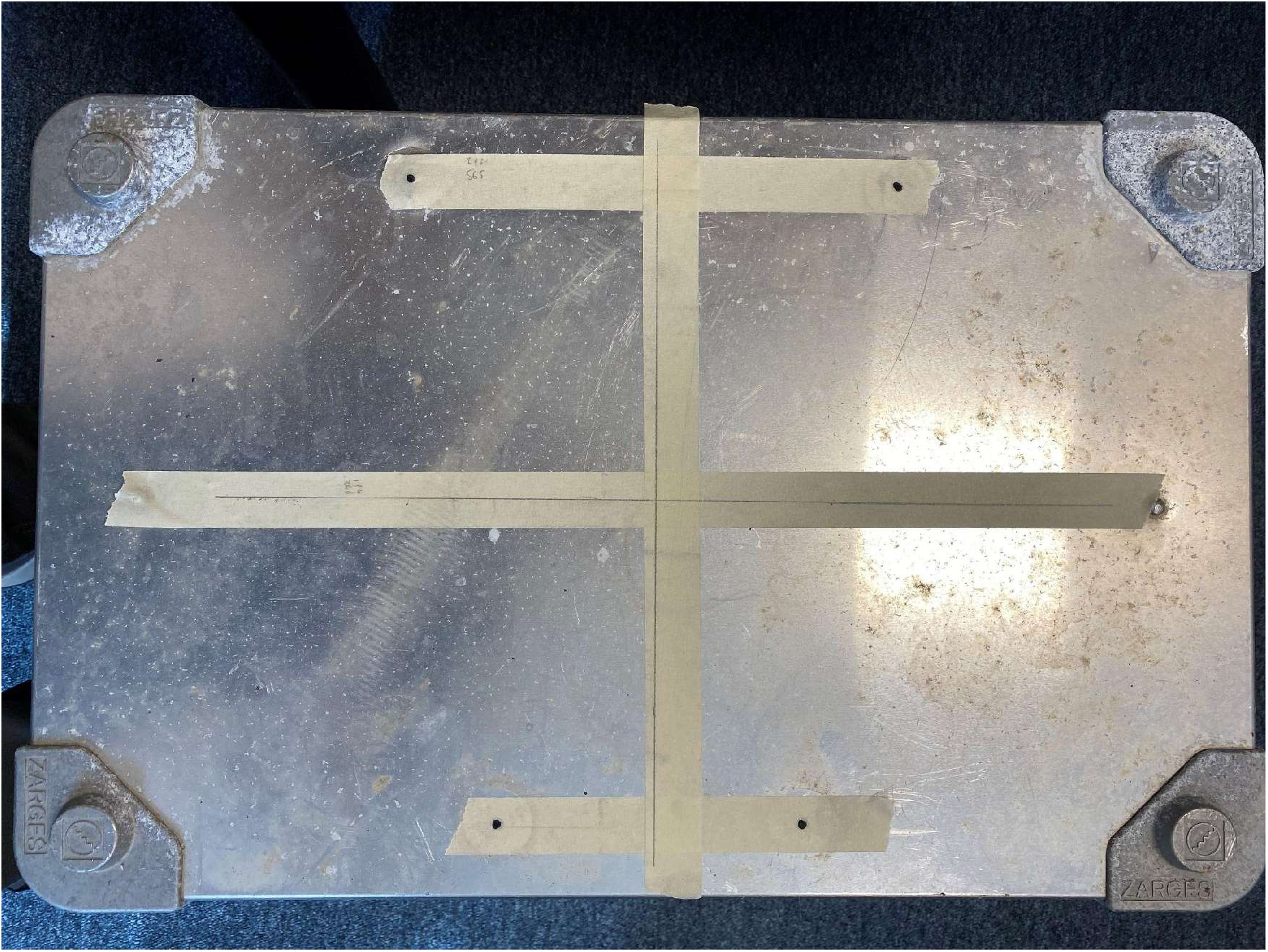
Holes drilled into the top of the battery box.

**Figure 5:**
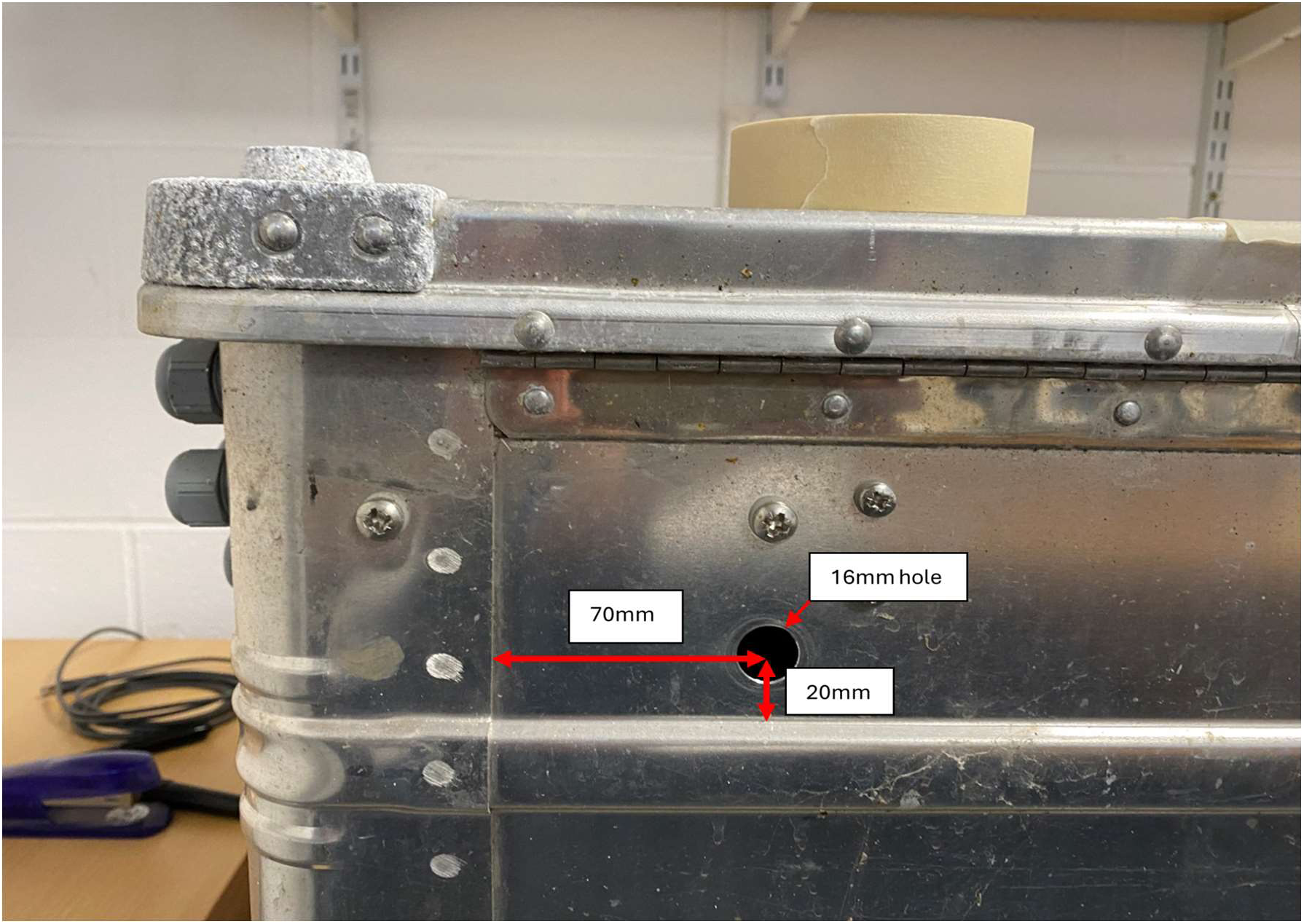
Marking up the hole on the back of the battery box.

### Assembling the electronics enclosure

The electronics enclosure is an IP67 enclosure that contains all the components that enable the central functionality through software operation and data capture. Figure 6 gives an overview of the electronics operating the UKCEH AMI-system, much of which is housed in the electronics enclosure. For detailed step-by-step instructions on installing components into the electric’s enclosure see the build manual [12]. The procedure is outlined here in summary.

1. Install the DIN rail into the electronics enclosure and clip on the terminal blocks and the timer relay (figure 7).
2. Secure 12 V regulator mounting plate to the enclosure box using two polyfix 4 x 10 mm screws and mount the 12 V regulator with two number 4 self-tapping screws (figure 8)
3. Mount the Raspberry Pi on top of the DINrPlate and then mount the watchdog on top of the Raspberry Pi on stand-offs (figure 9).
4. Mount the header extender on the RTC, and mount them onto the watchdog
5. Make sure the battery (CR 1220) is inserted into the Real time clock, this should last for 5 years with an accuracy of +/- 1 minute per year.
6. Plug the watchdog into the 12 V to 5 V stepdown and mount the assemblage of Raspberry Pi, watchdog, RTC and DINrPlate onto the DIN rail above the timer relay (figure 10)
7. The camera and lighting cables come out of the end of the protective conduit from the back of the camera box and can be seen in figure 11. To thread and connect the cables into the electronics box unscrew and remove the thin black plastic conduit connector nut (step 1 in figure 11)
8. Thread the USB and power cables through the hole in the electronics box (step 2 in figure 12)
9. Push the conduit and connector as far into the hole (step 2 in figure 12) as it will go, then replace the nut removed in step 1 (figure 11), passing it over the USB and power cables, then screwing it onto the conduit connecting until tight, this should be slightly more than hand tight so use a spanner. (step 3 in figure 13)
10. Connect the USB into the port shown in figure 12, step 4.
11. Connect the incoming 12 V supply to the positive and negative on the timer relay. Then link the positive to terminal 18 on the 12 V timer relay. Finally take a link wire from the negative on the timer relay to the black terminal block. Figure 14 shows, in more detail how the timer relay should be wired up.
12. Wire the other connections in the terminal block to the corresponding components as shown in figure 15.
13. Finally wire up the 12 V regulator as shown in figure 16

**Figure 6.**
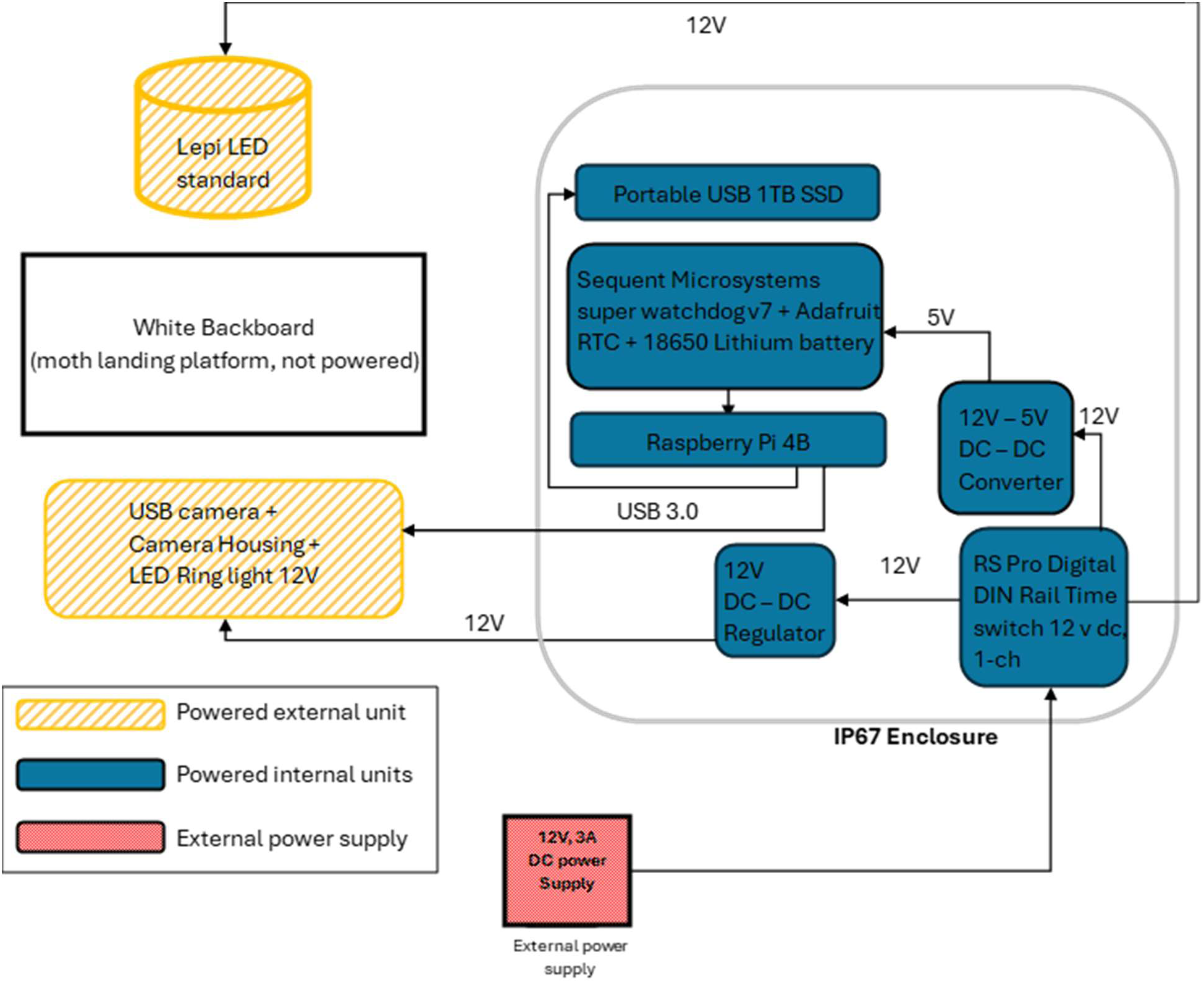
A schematic diagram for the electronics operating the UKCEH AMI-system, most of this is housed inside the electronics box (grey line).

**Figure 7.**
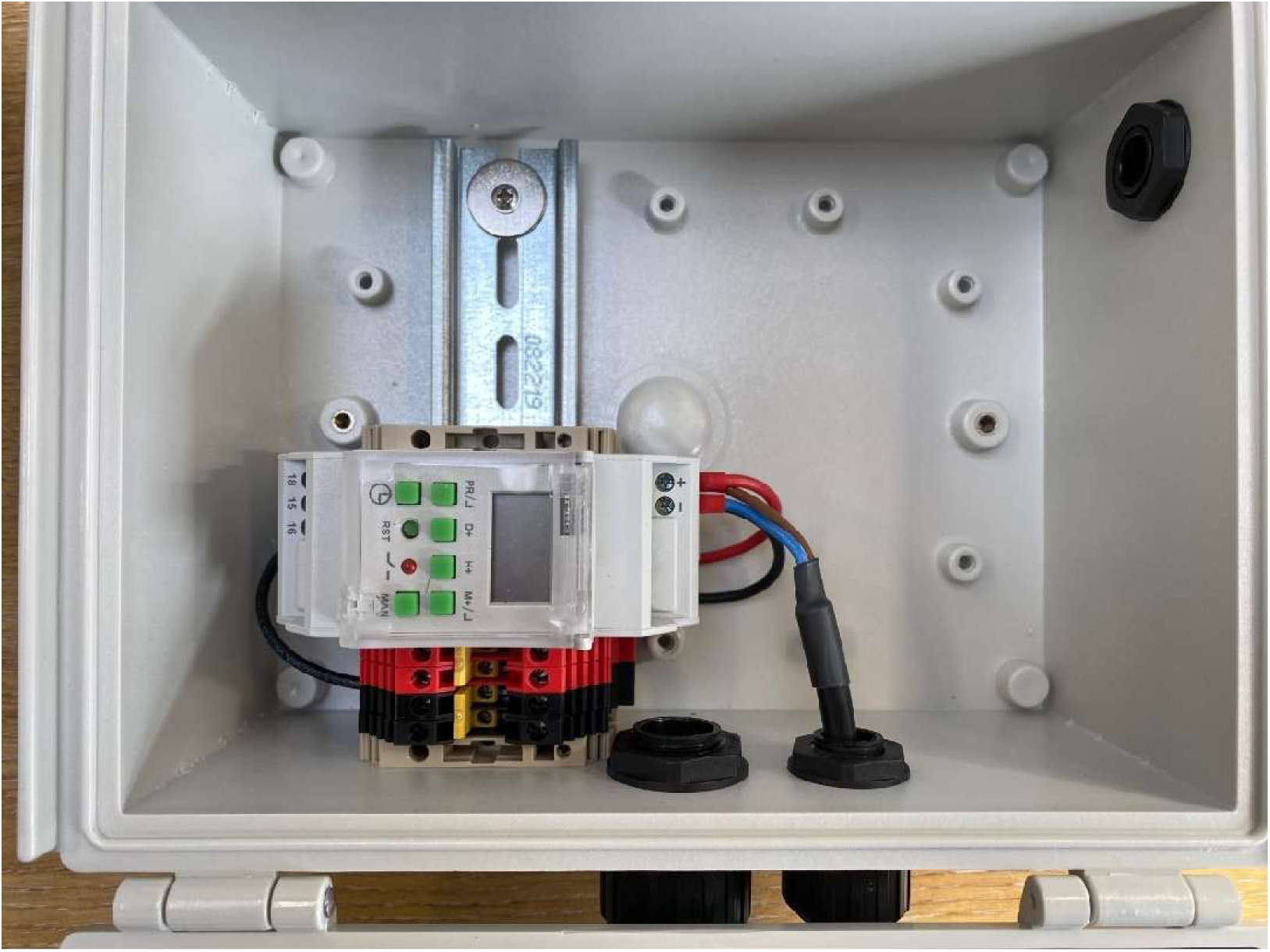
Terminal block and timer relay installed on DIN rail (wiring described in later steps)

**Figure 8.**
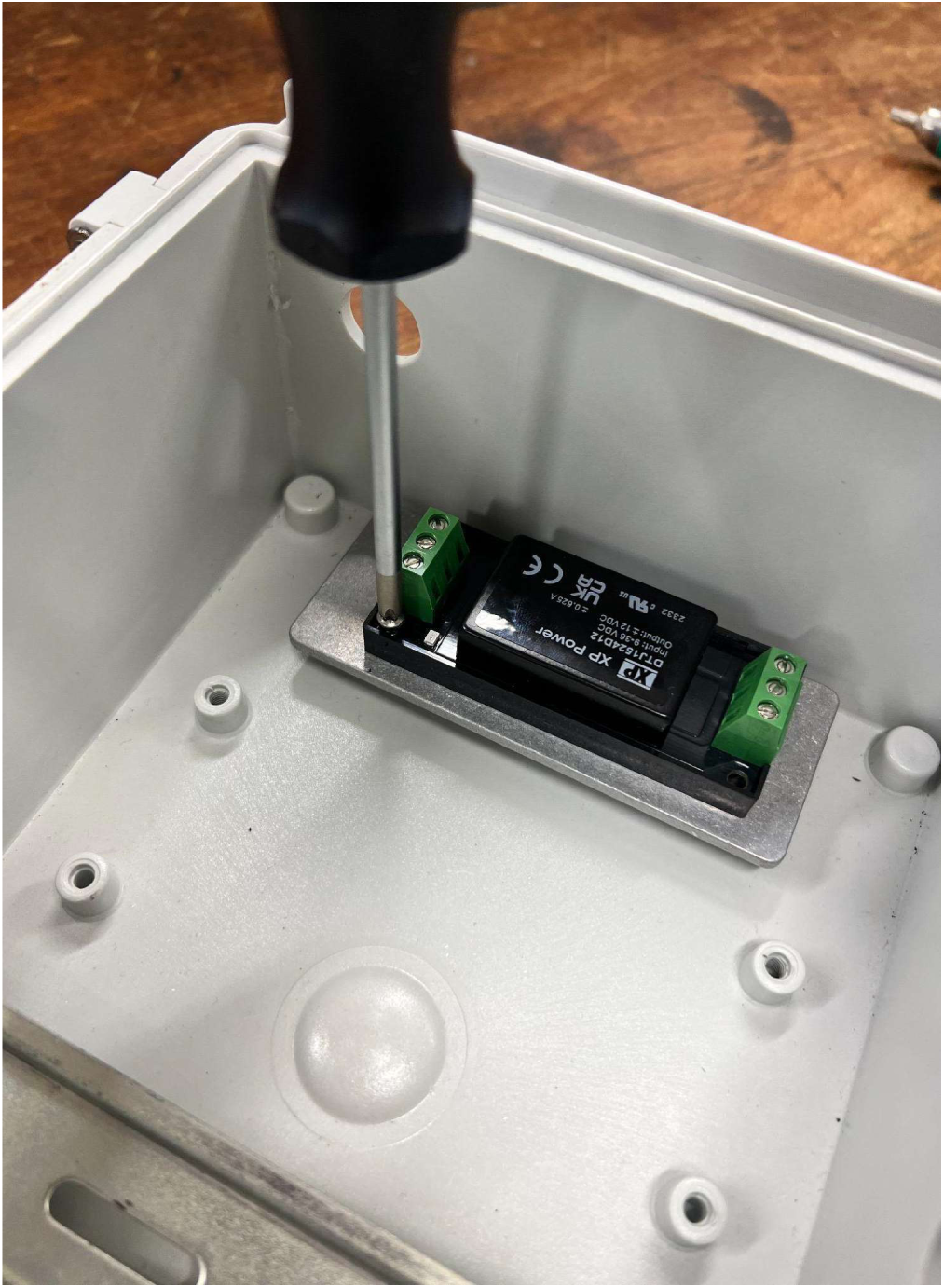
Mounting the 12 V regulator.

**Figure 9.**
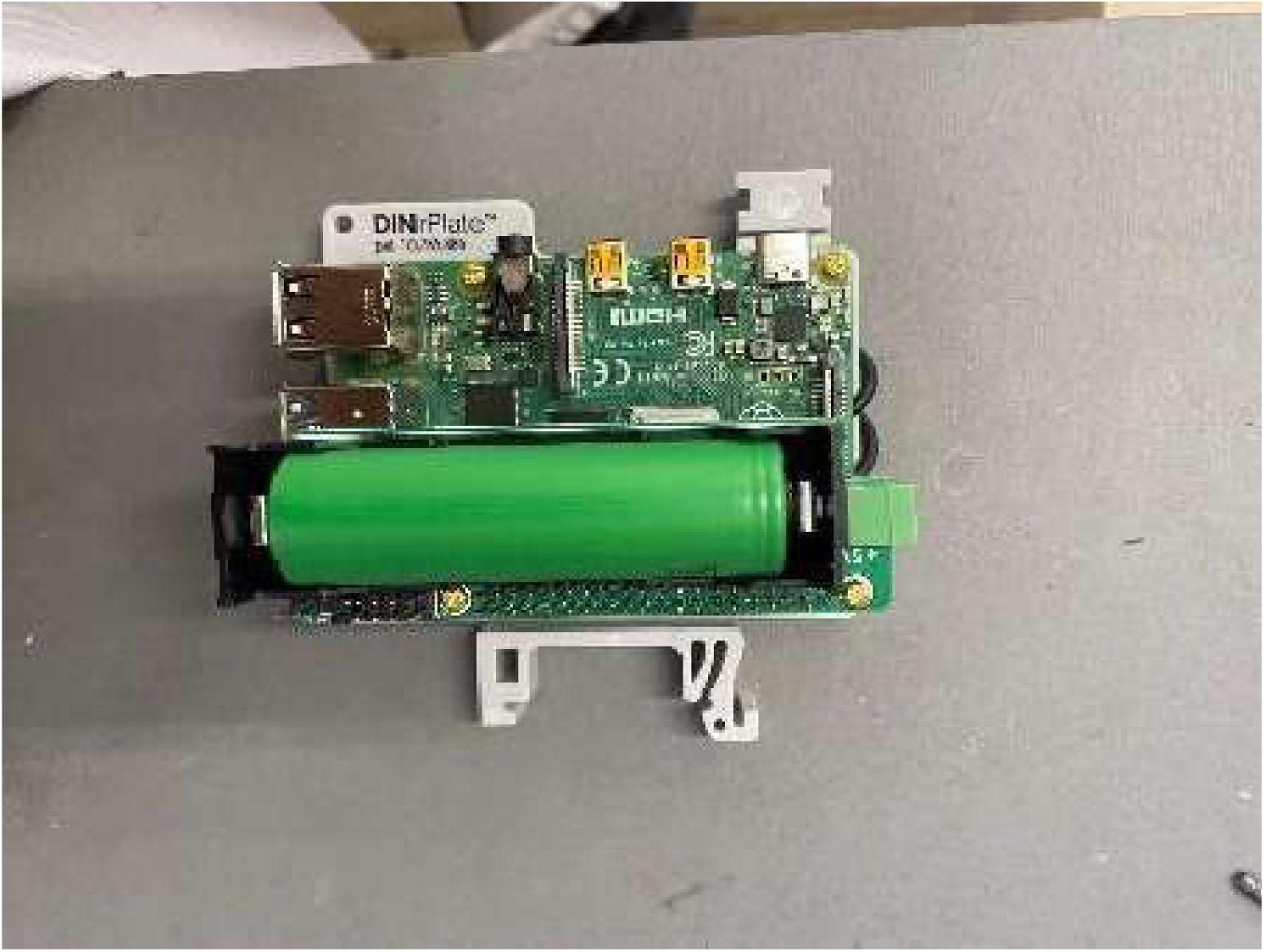
Raspberry Pi and Watchdog mounted on top of DINrPlate.

**Figure 10.**
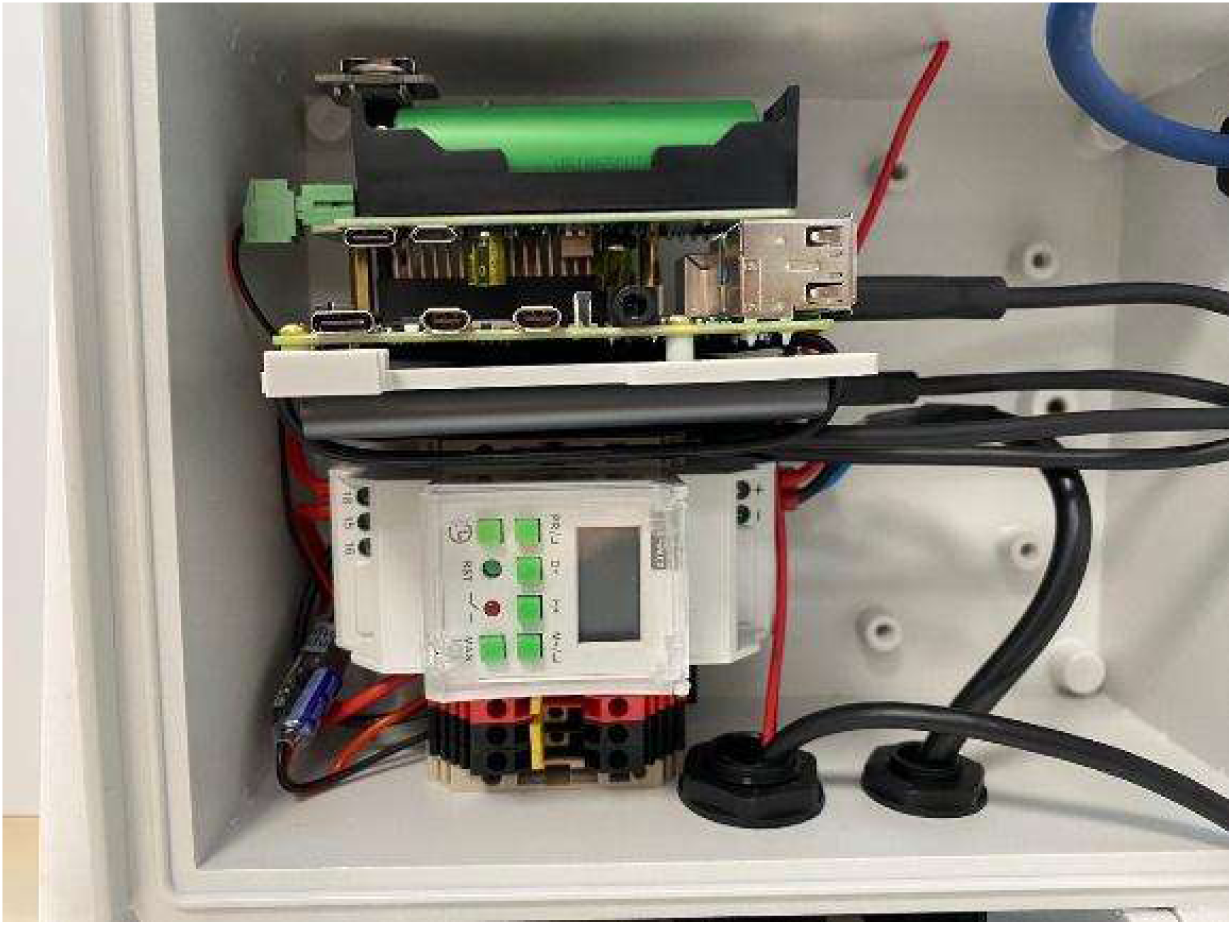
All components attached to the DIN rail (wiring described in later steps)

**Figure 11.**
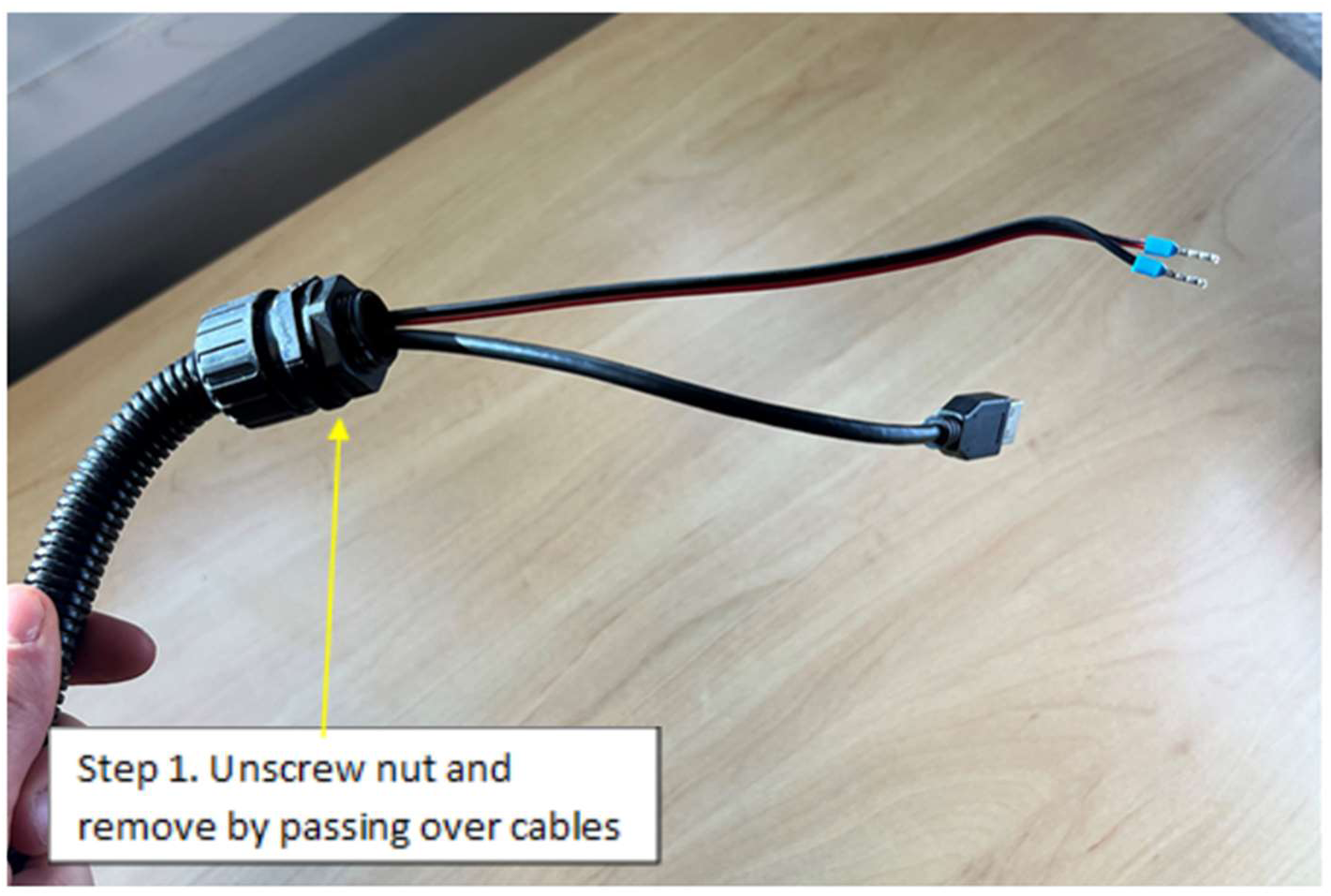
Conduit nut is first removed from cables extending from the back of the camera housing.

**Figure 12.**
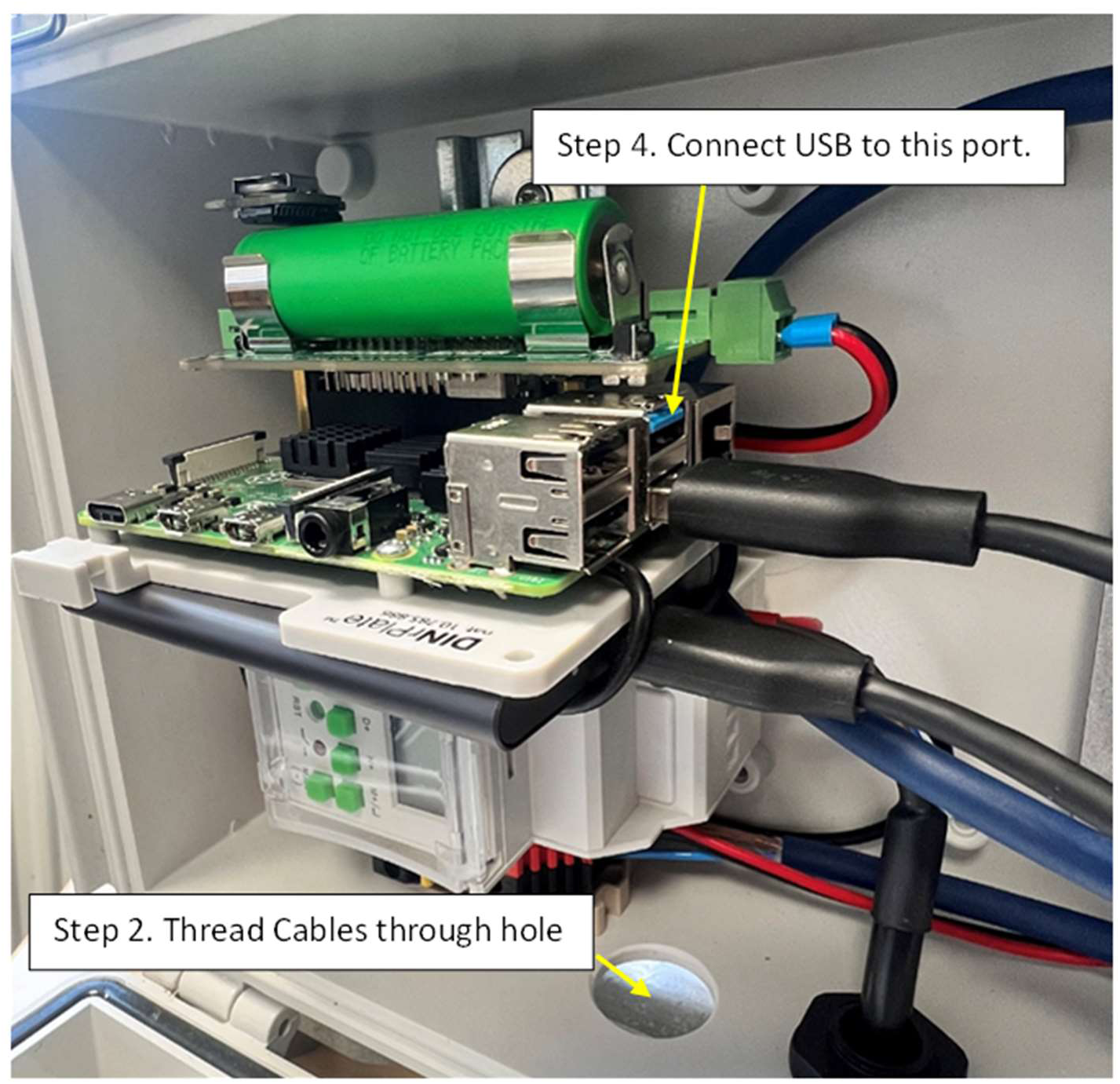
Wiring the camera and lighting ring into the electronics enclosure, steps 2 and 4.

**Figure 13.**
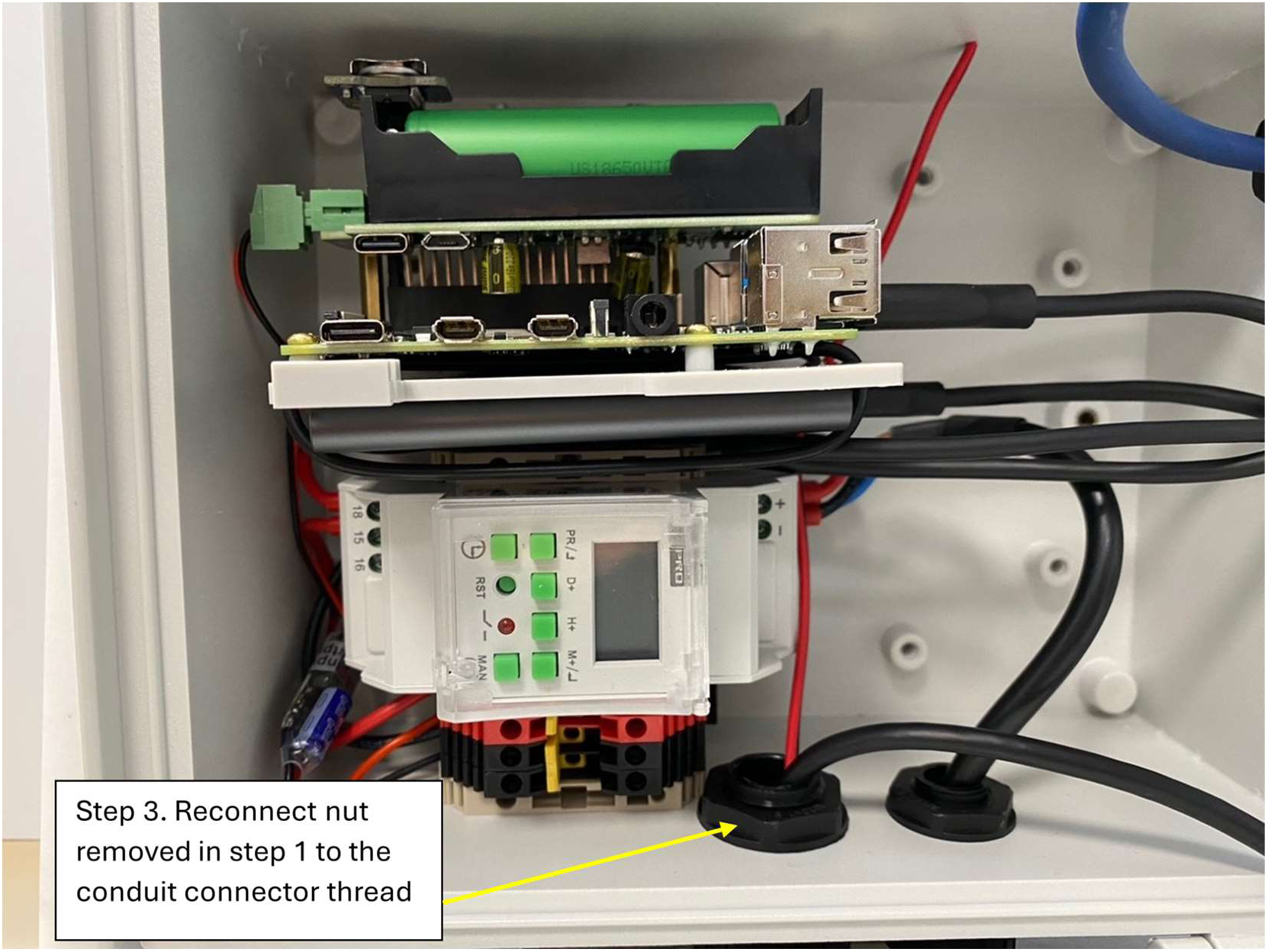
Wiring the camera and lighting ring into the electronics enclosure, step 3.

**Figure 14.**
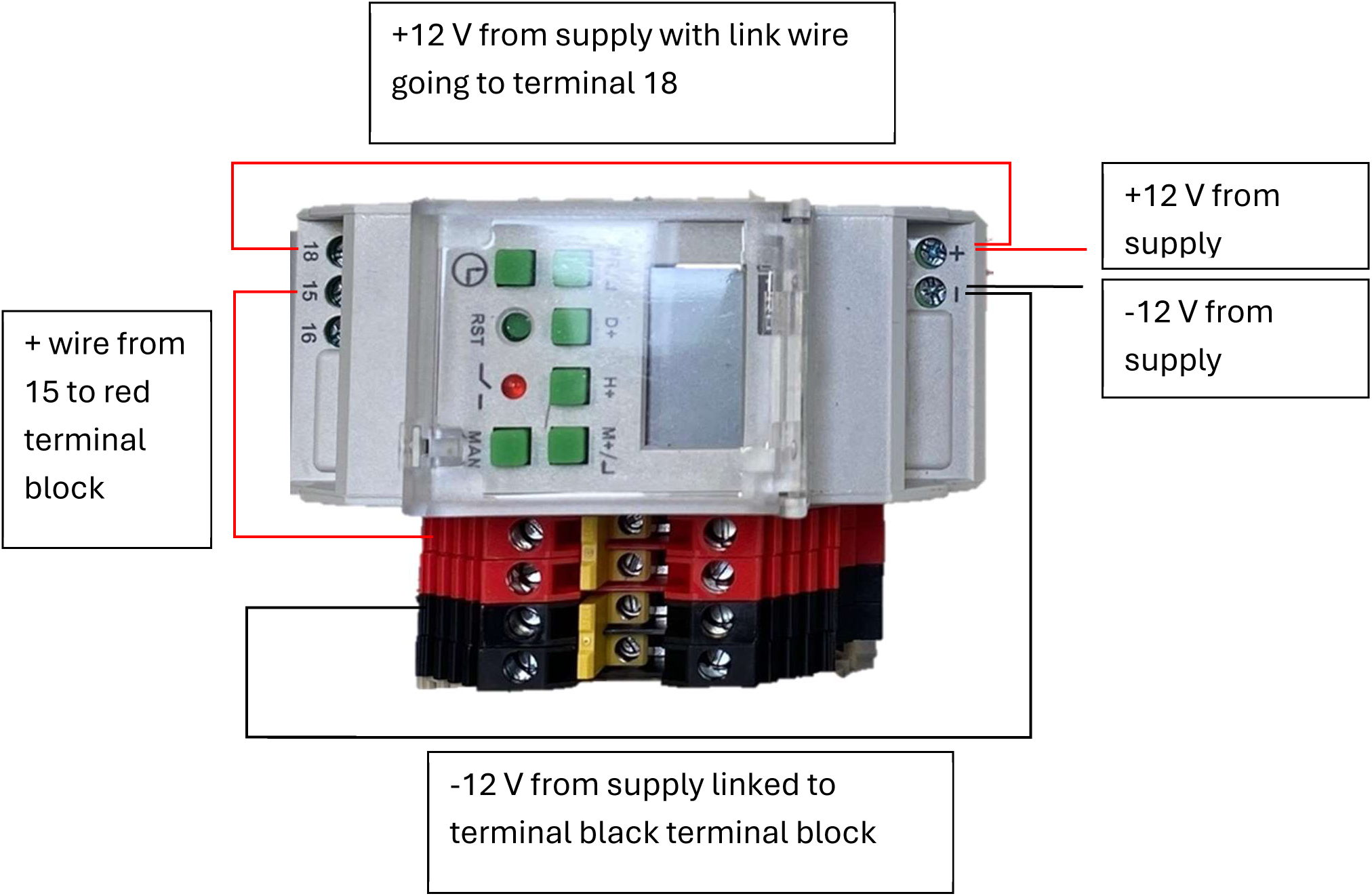
A schematic of the wiring of the timer relay, including connections from the 12 V power supply and connections to the terminal block.

**Figure 15.**
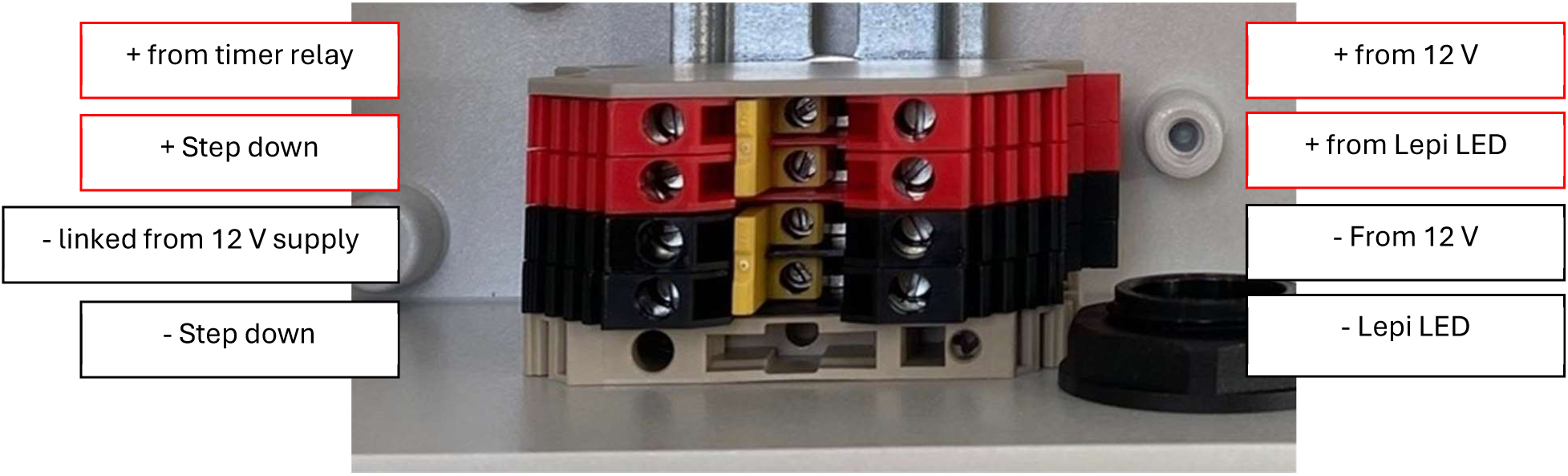
A schematic of the wiring into the terminal block.

**Figure 16.**
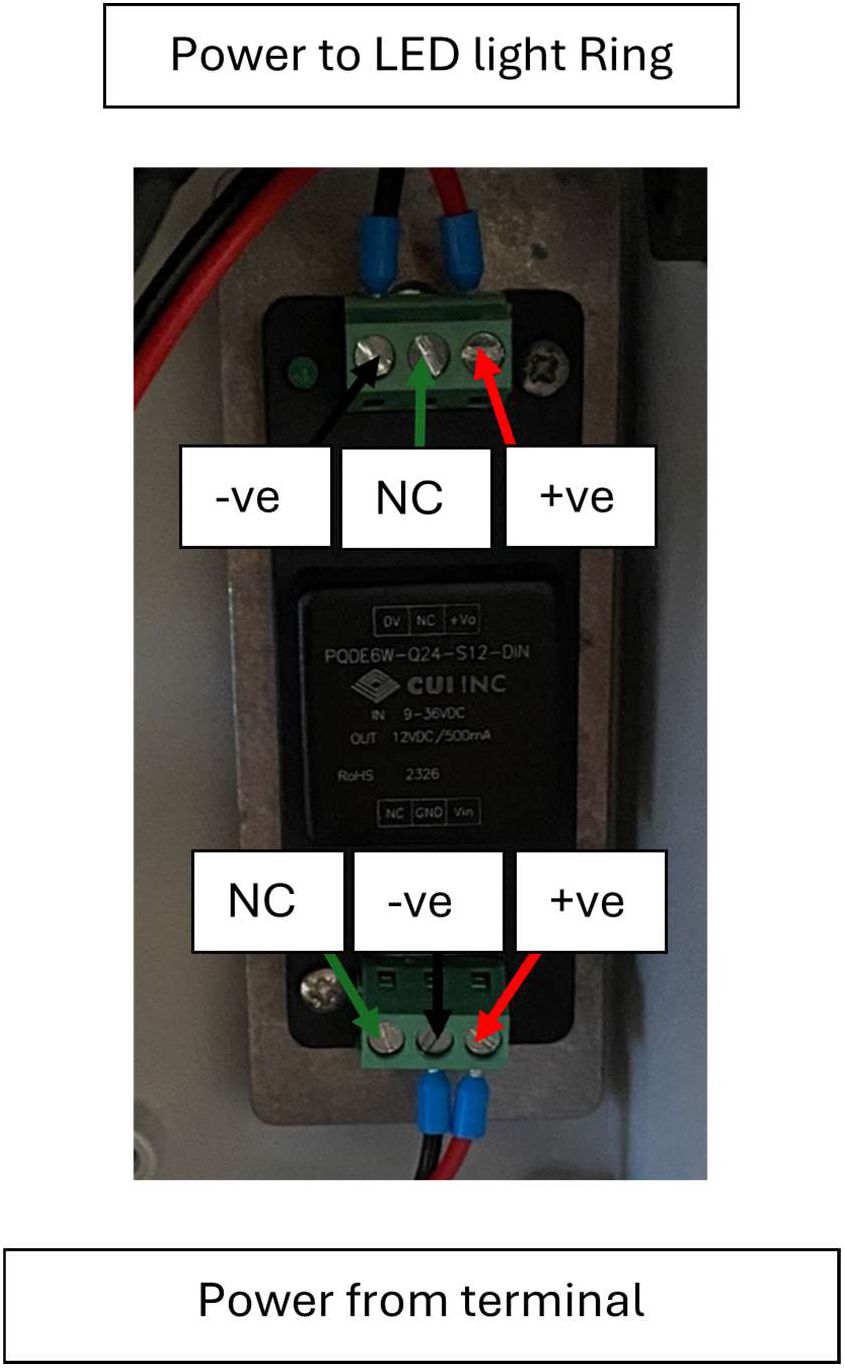
Wiring schematic for the 12 V regulator.

The fully assembled electronics enclosure is shown in figure 17.

**Figure 17.**
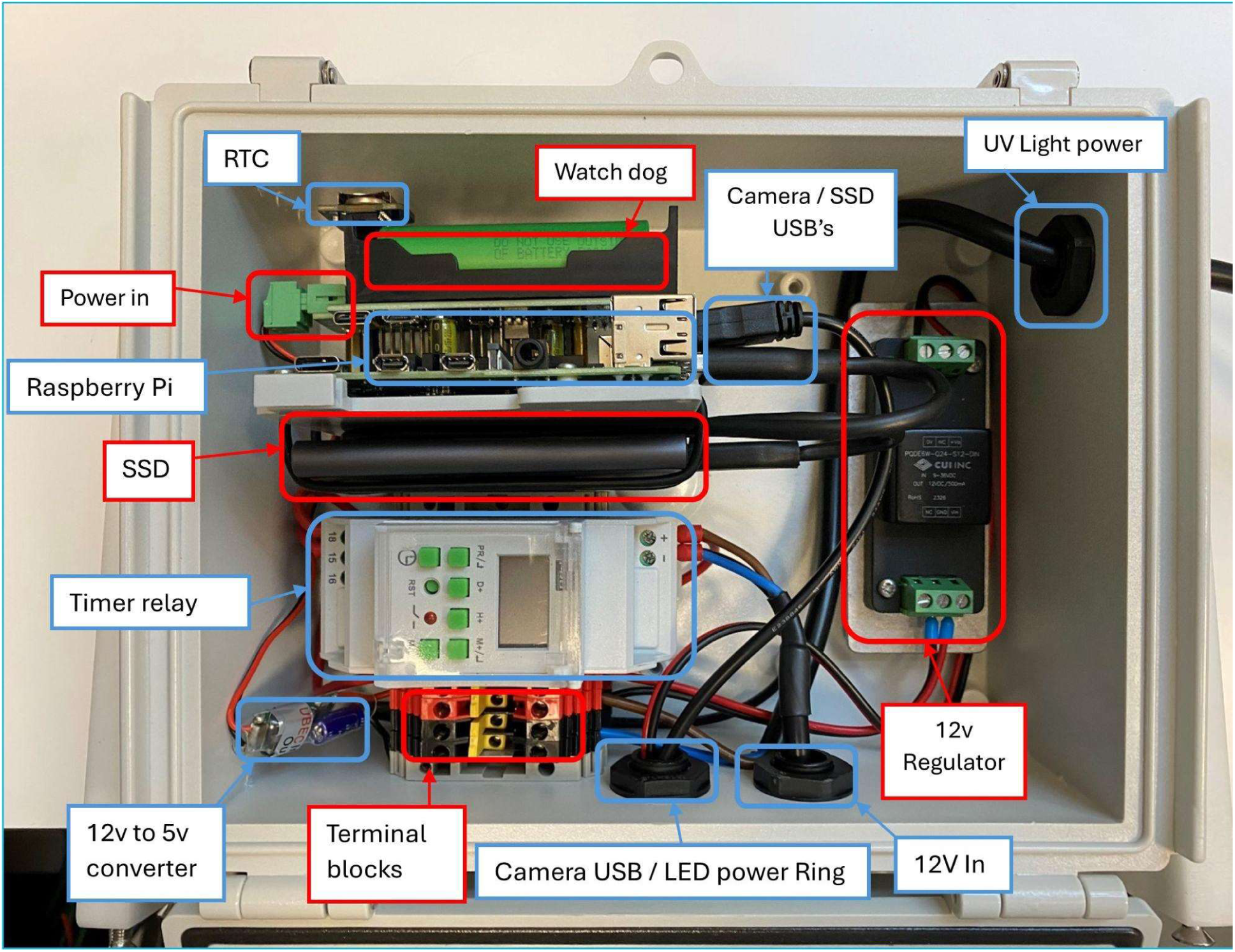
Fully assembled electronics enclosure.

### Assembling the backboard

Mount the back board to the uprights using four 25 mm M5 hex countersunk bolts ensuring the spacers are between the upright and back board. Then mount the electronics enclosure to the back board using four poly fix 5 x 10 mm screws (figure 18).

**Figure 18.**
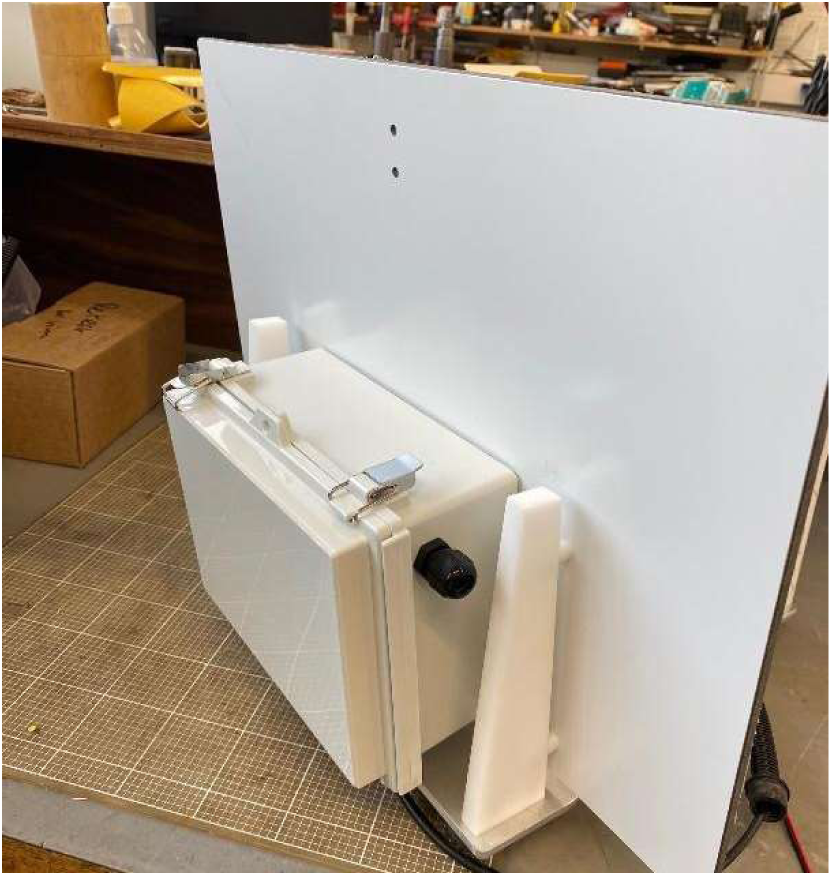
Back board attached to the electronics enclosure and uprights.

### Assembling the baseplate

1. Cut out and drill holes in the base plate as described in the drawing ‘DRAW3 Base plate V1.0.pdf’ [12].
2. Attach four standoffs (DRAW10 stand off V1.0.pdf) using four M5 x 16 mm countersunk socket screws (figure 19).

### Assembling the system

1. Once all necessary parts are prepared, the back board assembly and camera housing are mounted to the base plate, as shown in figure 20.
2. The conduit carrying cables from the back of the camera enclosure to the electronics enclosure is routed under the baseplate, from front to back and between the standoffs (figure 21)
3. The Lepi LED is then mounted to the back board using the UV light bracket as shown in figure 22. Finally, mount the UKCEH AMI-system (figure 23) to the battery box lid.

## 6. Operation instructions

Please be aware once the instrument is connected to a power supply, if the white lights come on this will mean the UV light is also on. This contains ultraviolet rays and can be harmful to look at. Please take extra caution, covering the lamp or wearing UV protective glasses. The light can be de-activated by disconnecting it, but in normal conditions the UV light will always be on when the system is on.

**Figure 19:**
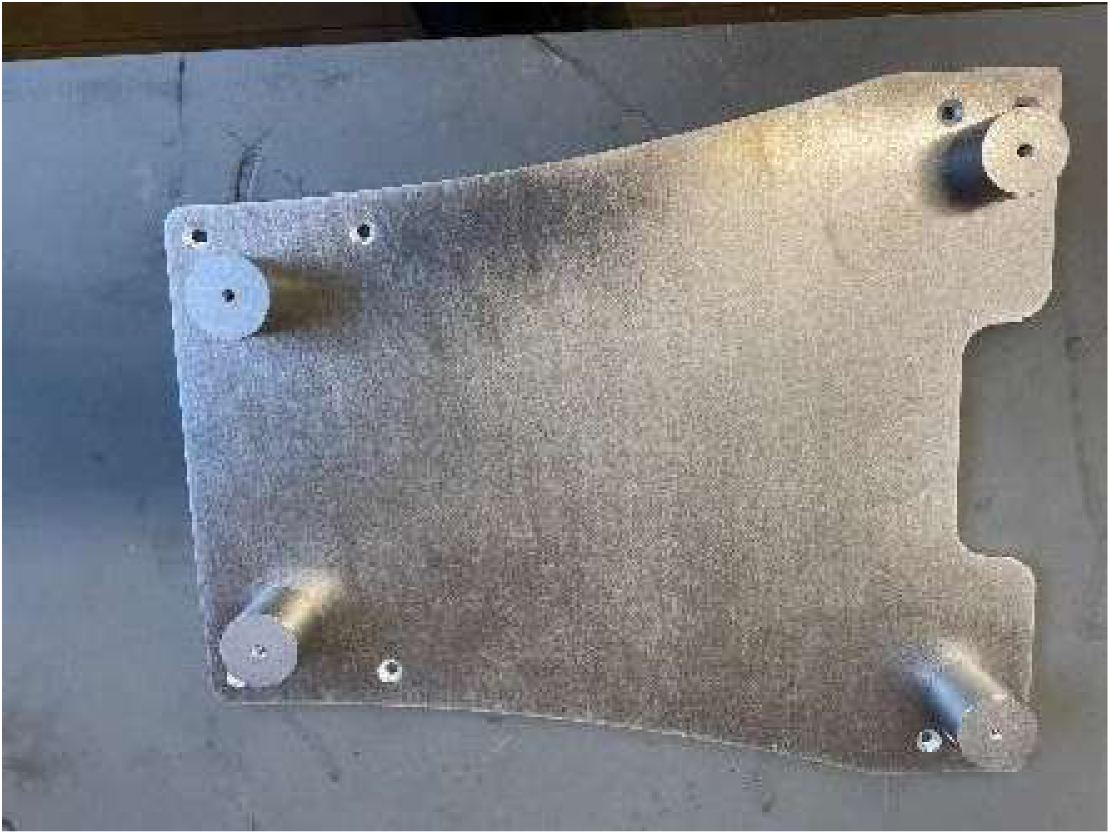
Standoffs attached to the underside of the baseplate.

**Figure 20.**
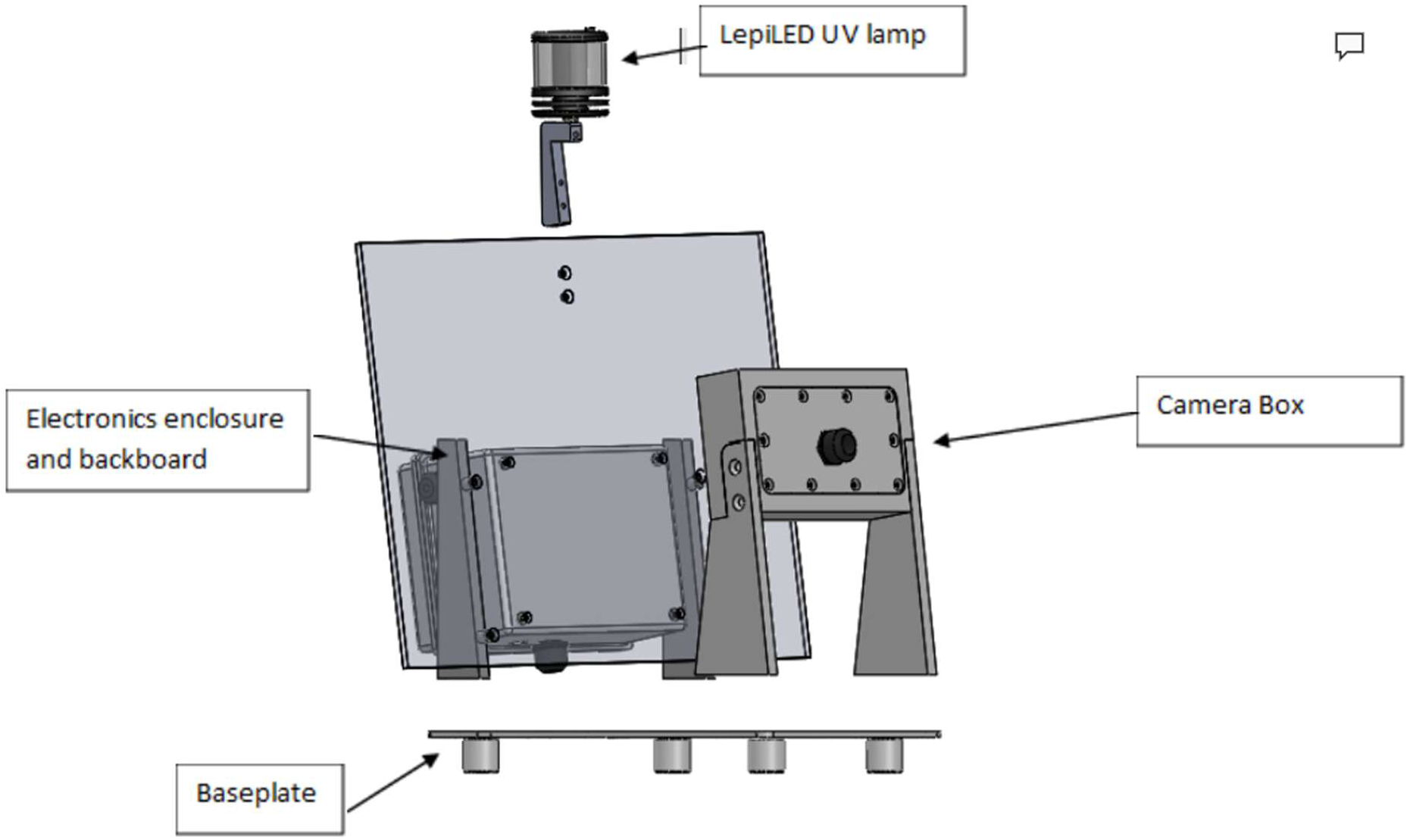
Assembling the UKCEH AMI-system.

**Figure 21.**
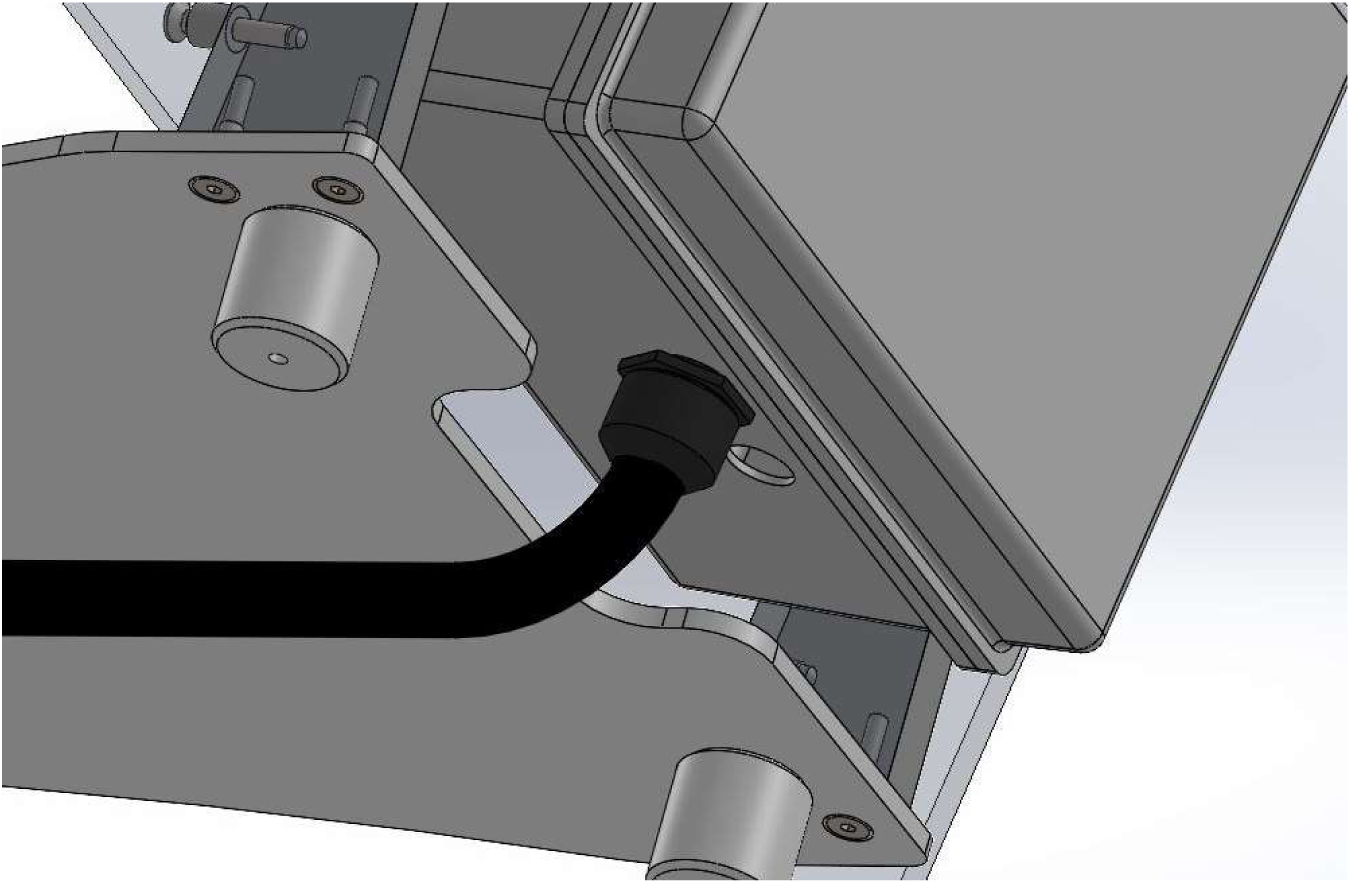
Cable routing from the camera housing to the electronics enclosure.

**Figure 22.**
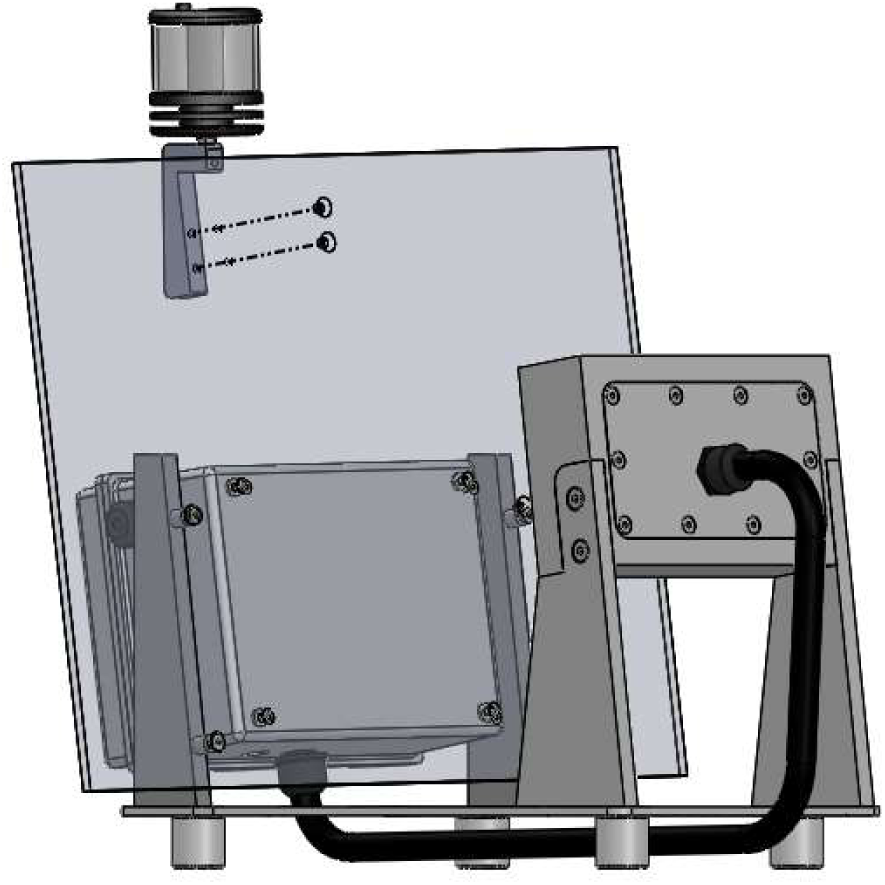
Attaching the Lepi LED.

**Figure 23:**
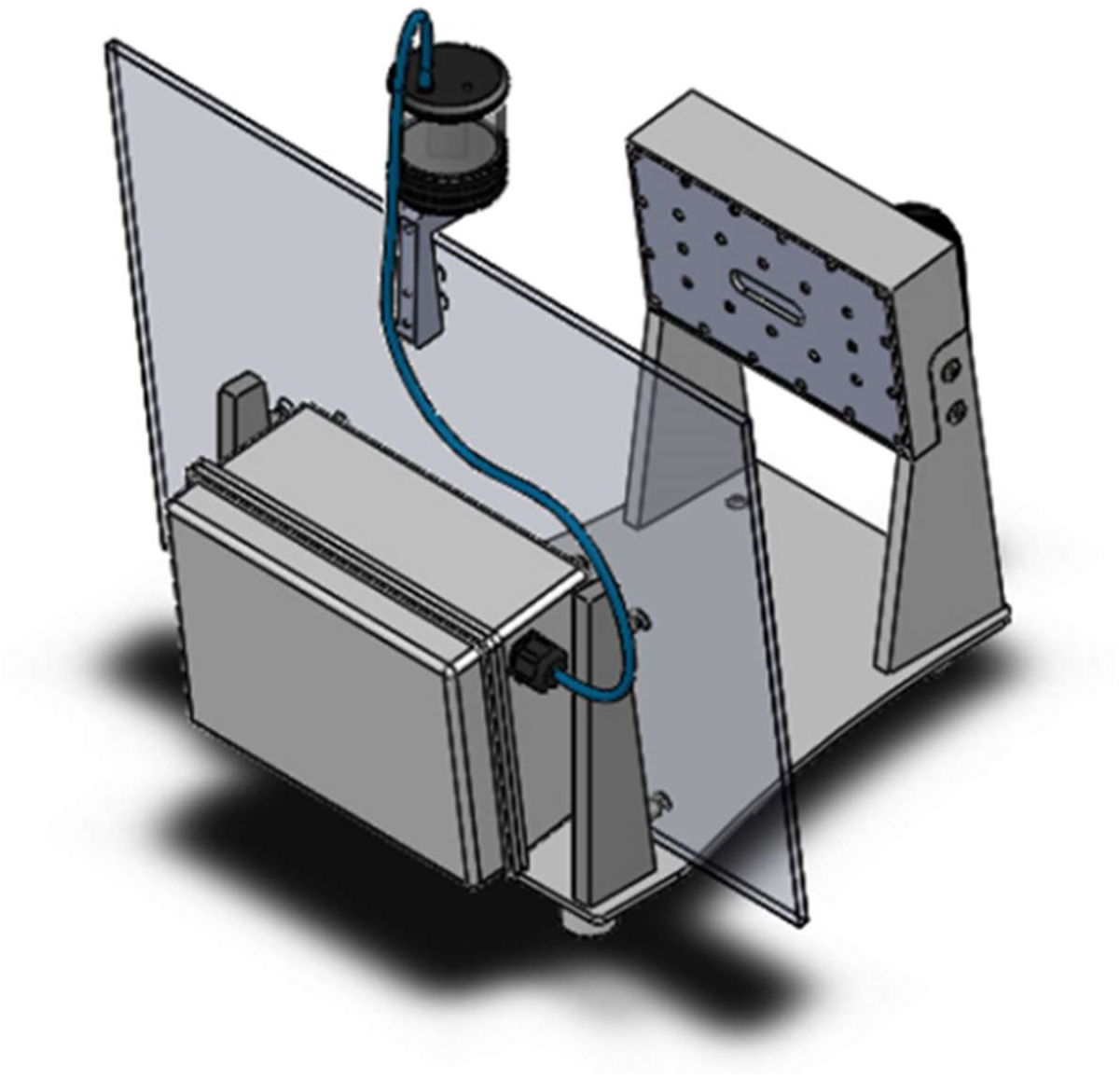
Final assembly view.

Prior to operation, the UKCEH AMI-system needs configurations and settings as below:

Flash the Raspberry Pi OS image [12] onto the 64 GB SD card. This image has the required software for operating the system installed. To connect to the systems, plug a computer into the RaspberryPi using an ethernet cable and use SSH. The address for the Raspberry Pi is and the default password “Nature17”. Change the password after first connecting to ensure security.

### Setting time on the real time clock

The most effective way of setting time on the RTC is to connect the Pi to the internet, doing so will sync the Pi’s time to an NTP (Network Time Protocol) server automatically. If an internet connection is not available the following method can be used.

1. Open the terminal on the Raspberry pi and check the time with the command ‘timedatectl’
2. If necessary update the time zone, for example ‘sudo timedatectl set-timezone Europe/London’

a. You can see a list of timezones using ‘sudo timedatectl list-timezones’
3. Check the timezone has been updated by running ‘timedatectl’
4. Update the date and time to the current time e.g. ‘sudo date -s “2025-12-30 14:52”
5. Verify the date and time by running ‘date’
6. Then you need to write the time into the RTC battery with the following command ‘sudo hwclock -w’
7. Confirm the RTC battery time is correct with ‘sudo hwclock -r’

### Testing the UKCEH AMI-system

1. Set up and configure the timer relay following the manufacturer’s instructions. The 12 V timer relay is procured directly from RS Components, part number 8966891
2. Using the timer relay, turn the system ON
3. Cover the UV light/LepiLED to protect yourself from the UV light. This is a precautionary measure but is recommended if working close to the light for extended periods of time.
4. Once the AMI is ON it will begin to take one image every ten seconds.
5. After a minute use the timer relay to turn off the system.
6. Once the AMI is OFF, wait until the watchdog finishes switching OFF the raspberry pi, this will be apparent when all of the LEDs on the Raspberry Pi turn off.
7. Take the SSD out and connect it to your computer to see if the images have been taken properly and the filenames have the correct time.

### Solar panel and battery recommendations

For field deployments requiring long-term autonomous operation, we recommend the following solar panels and batteries, to be used with a Sunsaver 20L solar charge controller:

- 2 x 100 W solar panels in parallel (12 V panels)

- Max open circuit voltage (Voc): 25 V
- Max short circuit current (Isc): 25 A
- 2 x AGM Lead Acid deep cycle / marine batteries in parallel (12 V)

- 120 AH 12 V Absorbed Glass Mat (AGM) or Marine Lead Acid Battery

This would support an operational program turning the instrument on for 10 hours a night every 3^rd^ night (tested at latitude 52 degrees North), however additional batteries and panels will be required in areas that are shaded, or experience limited sunlight, such as in high latitudes.

## 7. Validation and characterization

The UKCEH AMI-system is designed to operate in tropical and temperate climates, which requires the system to be resistant to high humidity and rain, as well as heat. To test the system’s resistance to heat the system was tested by running at a high temperature with a maximum load on the processor over 16 hours (figure 23). The camera housing and electronics enclosure were run in a climate-controlled environment and based on the results of this test we are confident that the UKCEH AMI-system can operate in high temperatures without impact on the electronics.

**Figure 23.**
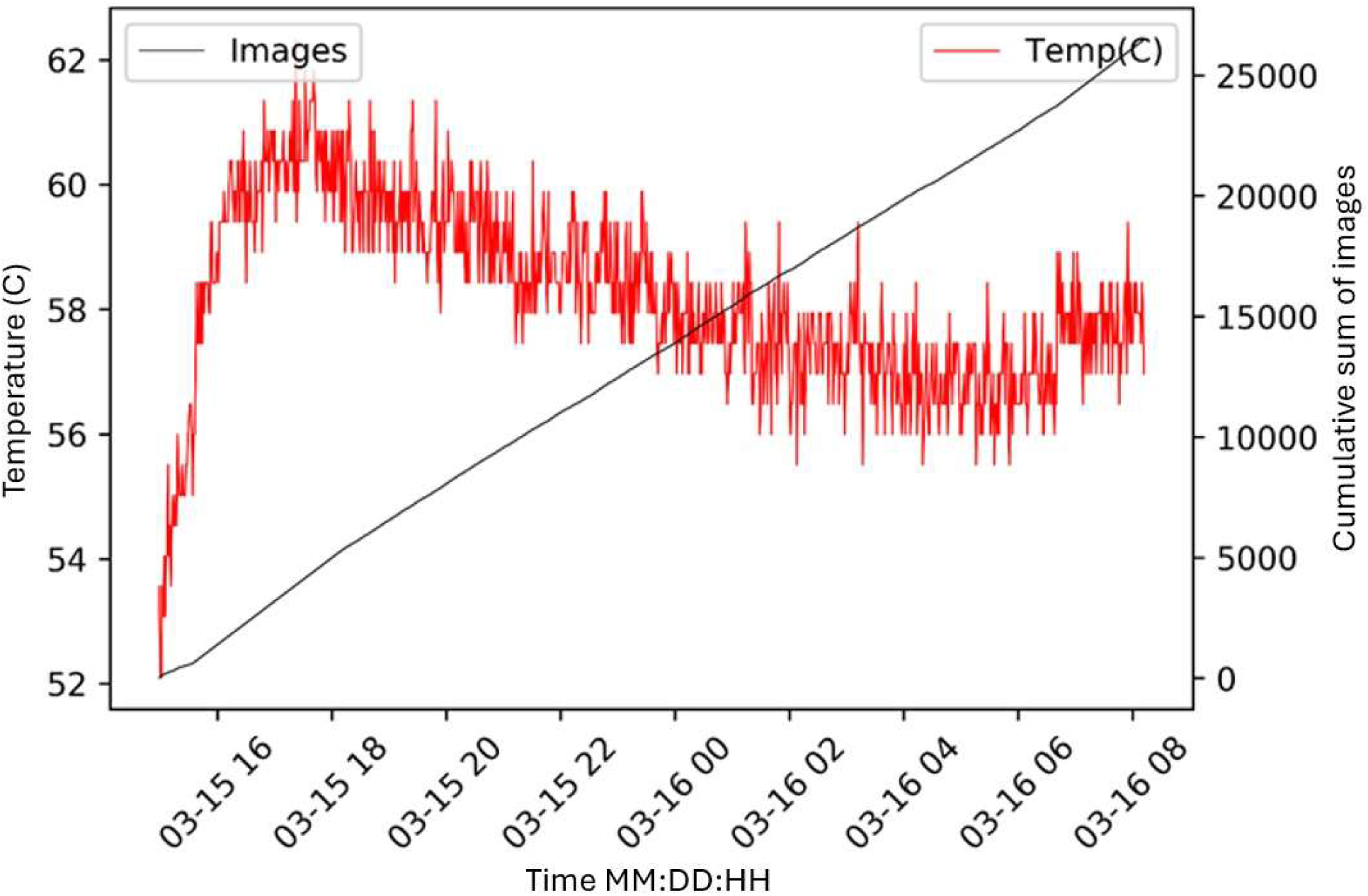
Results of running the UKCEH AMI-system through a heat test shows that the system continued to capture image continuously in temperatures up to 62C over a period of 16 hours.

To ensure the camera housing was watertight we submerged it in water for 30 minutes at a depth of one meter, replicating an IPx7 waterproof test. During this test no water entered the camera housing.

Whilst all the components used in the UKCEH AMI-system are electrical safety compliant, we additionally tested the system as a whole to ensure the electrical safety of the device. Continuity testing of the circuitry and EMC testing through an electrical compliance test house identified no electronics issues (figure 24).

**Figure 24.**
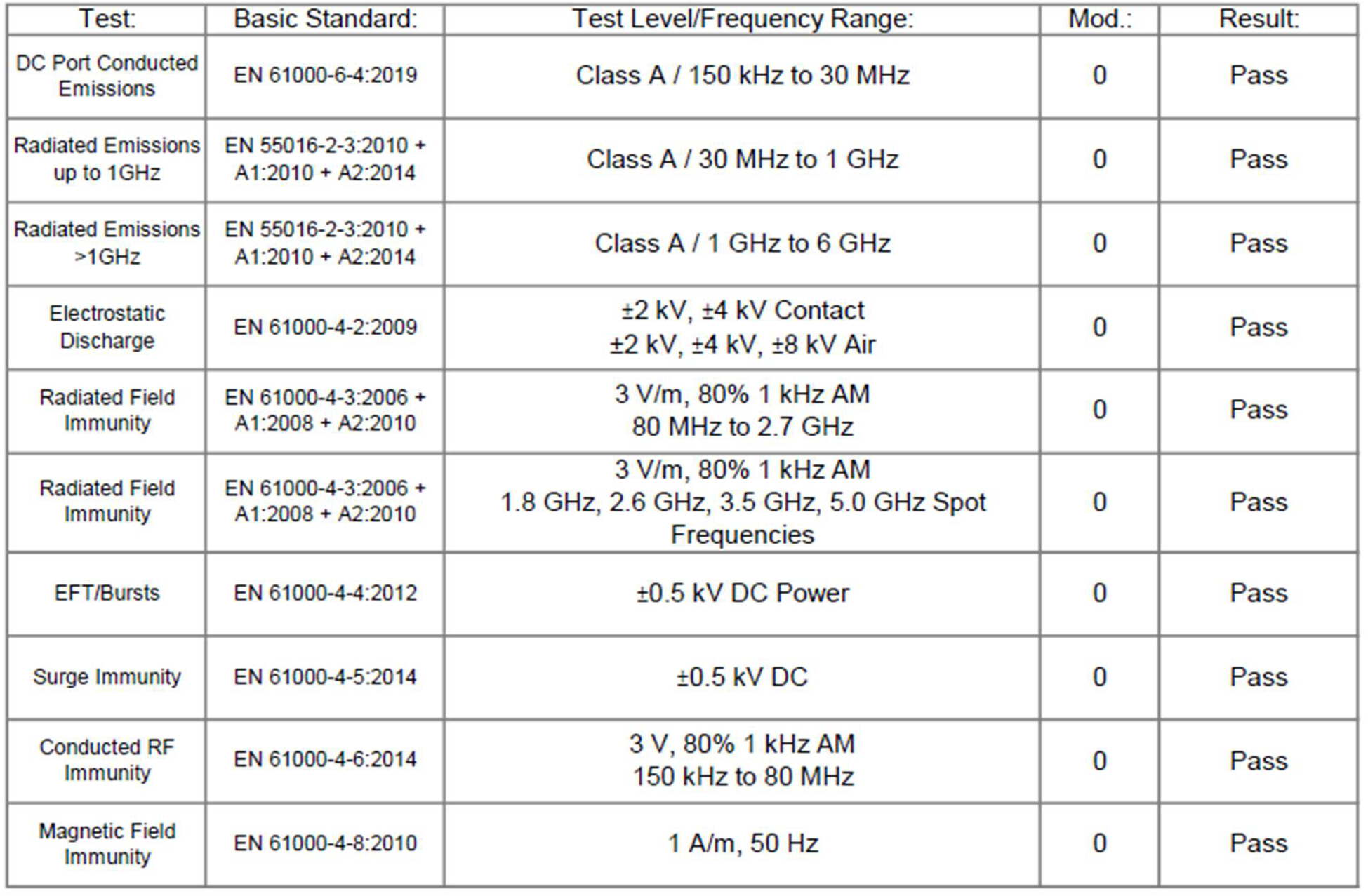
Summary of electrical testing.

As previously mentioned, nearly 200 UKCEH AMI-systems are deployed globally. These systems have collected many thousands of pictures an example of which is shown in figure 25. In the United Kingdom we run the systems for one night in every three, with three batteries and two 100W solar panels. This setup is enough to power the system from early May, through to late September. With a 500 GB SD card we find this is able to hold approximately 340,000 pictures. Our standard schedule is to take one image every 60 seconds between sunset and one hour before sunrise, meaning that one SSD can last an entire field campaign (May-September). Given the number of units in use we have also been able to identify common issues that users should be aware of. After prolonged periods in storage we recommend checking and replacing the Lithium 18650 battery and the RTC CR 1220 battery. In some cases these have lost charge and caused problems with subsequent deployments. When closing the electronics enclosure, for example after swapping the SSD drives, it is important to check that cables are not trapped in the seal as this can allow water to enter the electronics enclosure. We recommend testing each system at the office/lab before deployment so that any issues can be identified before going to the field.

**Figure 25.**
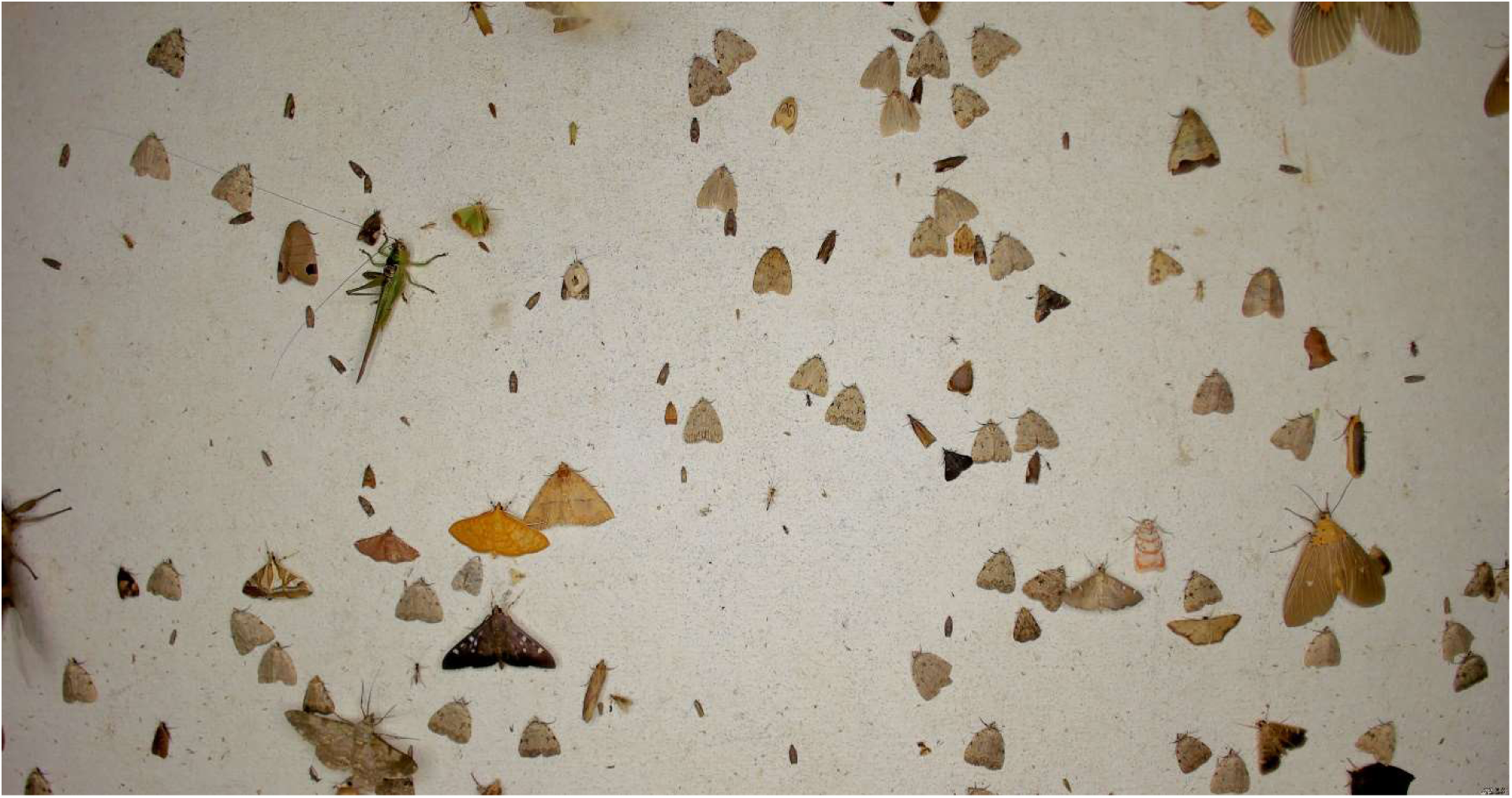
Example image from an UKCEH AMI-system deployed in Thailand. Images have a resolution of 4096 x 2160 pixels.

As evidence of the AMI functionality, we here show data from a network of systems that have been located at a number of sites in Cost Rica. The images from these systems have been analysed with computer vision tools to count all instances of insects on the backboard. The analysis of these data (figure 26) shows the variability in the number of detections made between days, sites, and across the seasons. An additional example of the use of the UKCEH AMI-systems can be found in Bett *et al.* 2025 [17], where the authors deploy an AMI system in Kenya, reporting an average of 10 insects per night over two months.

**Figure 26.**
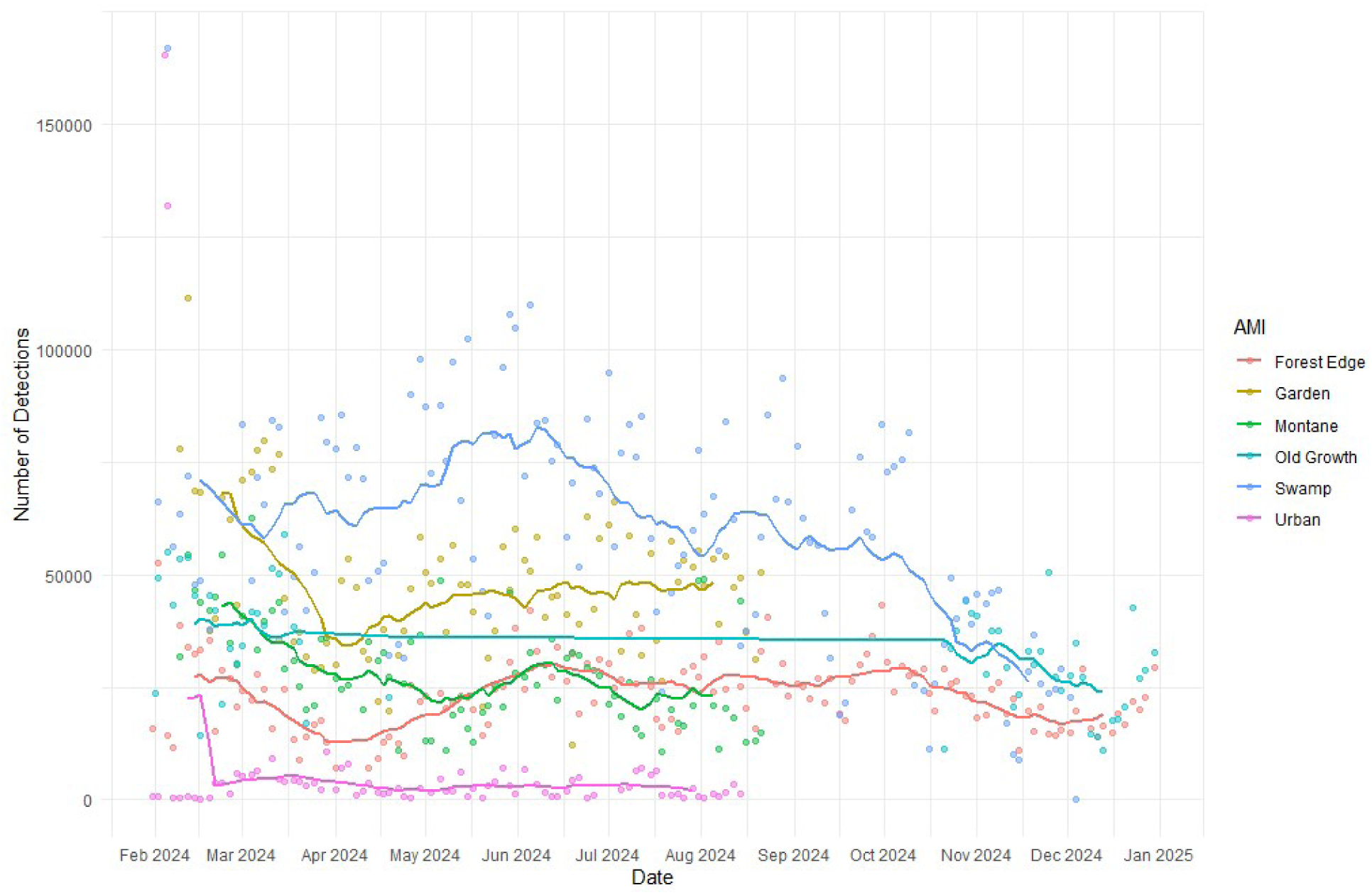
Plot of the number of moths detected each night across 6 UKCEH AMI-systems deployed in Costa Rica. Points give raw numbers per night and lines represent a 14-day moving average. In this analysis the same individual moth in multiple images will be counted as multiple detections. An AI classifier has been used to remove non-moth species from the data. The graphs demonstrate the variability we observe in the number of moths that come to the trap across nights, seasons and locations, offering the possibility to explore drivers of these trends, such as habitat, weather and climate. In this example it is notable that the urban site has very low number, as would be expected.

## CRediT author statement

**William Lord**: Conceptualization, Methodology, Investigation, Writing - Original Draft, Writing - Review & Editing. **Simon Teagle**: Conceptualization, Methodology, Investigation, Writing - Original Draft, Writing - Review & Editing. **Joshua Alton**: Methodology, Investigation. **Gabriel Bannister**: Methodology, Investigation. **Jonas Beuchert**: Methodology, Validation, Writing - Review & Editing. **Kim Bjerge**: Software, Conceptualization, Methodology, Investigation, Writing - Review & Editing. **Dylan Carbone**: Validation, Formal Analysis, Investigation. **Alba Gomez Segura**: Software, Methodology, Validation, Investigation. **Tim Howson:** Methodology, Investigation. **Toke Thomas Høye**: Conceptualization, Funding acquisition, Methodology, Writing - Review & Editing. **Jenna Lawson**: Validation, Formal Analysis, Investigation. **Abhi Ravivarma**: Methodology, Validation, Investigation, Writing - Original Draft, Writing - Review & Editing. **David B. Roy**: Funding acquisition, Validation, Writing - Review & Editing. **Dan Rylett**: Conceptualization, Methodology, Writing - Review & Editing. **Grace Skinner**: Validation, Formal Analysis, Investigation. **Alan Warwick**: Methodology. **Tom August**: Conceptualization, Funding acquisition, Validation, Investigation, Writing - Original Draft, Writing - Review & Editing

## Acknowledgements

We thank our partners from all over the world who have tested UKCEH AMI-systems and provided us with feedback to help to improve designs. We thank University of Embu, National Museum of Kenya, Tropical Biological Association, Dedan Kimathi University of Technology, Perivoli Rangeland Institute, Osa Conservation, Organization of Tropical Studies, Nanyang Technical University, National Science Museum Thailand, Chuo University, Smithsonian Tropical Research Institute, Makerere University, Jos Institute, Madagascar Biodiversity Centre, Mila - Quebec Artificial Intelligence Institute, Montreal Space for Life - Insectarium, Université de Sherbrooke, Natural Resources Canada, Vermont Center for Ecostudies, The Alan Turing Institute, Université Laval, and others who have supported our work.

This work was supported by the AMBER project funded by the Aberdeen Group Charitable Foundation, and the INSPIRE project funded by the Aberdeen Group Charitable Trust [a registered charity in Scotland SC040877]; the European Union’s Horizon Europe Research and Innovation programme under the MAMBO project [grant agreement no.101060639], Easy RIDER: Real-time IDentification for Ecological Research and Monitoring’ funded under the Natural Environment Research Council (NERC) Global Partnerships Seedcorn Fund [NE/W004216/1], the Natural Environment Research Council (NERC) and the Biotechnology and Biological Sciences Research Council (BBSRC) joint research program AgZero+: Towards sustainable, climate-neutral farming [NE/W005050/1]; This article is based upon work from COST Action InsectAI, CA22129, supported by COST (European Cooperation in Science and Technology)

## Notes

https://zenodo.org/records/18131879

### Competing Interest Statement

The authors have declared no competing interest.

